# Pathway-wide base editing charts chemical-genetic interactions in MAPK signaling

**DOI:** 10.64898/2026.09.25.753639

**Authors:** James Woods, Calvin XiaoYang Hu, Dong Man Jang, Camille Freedman, Sarah Canarelli, Hui Si Kwok, Irtiza Iram, Michael Eck, Brian Liau

## Abstract

CRISPR base editor (BE) scanning enables sequence-level interrogation of proteins at scale in their native cellular and genomic contexts. This approach opens new opportunities to dissect cellular pathways, where signaling depends on coordinated interactions among multiple pathway proteins. We apply BE scanning to the RAS-RAF-MEK-ERK (MAPK) cascade, a central oncogenic pathway and major therapeutic target. Compounds targeting the MAPK pathway have mechanisms of action and resistance that remain incompletely characterized. By base editing 22 MAPK genes under either hyperactivation or inhibition at various nodes, we present a chemical-genetic map of the signaling pathway. Pathway-wide analysis shows that drug resistance mutations frequently occur in proteins other than the direct drug target itself, and we exploit these chemical-genetic interactions to recapitulate key pathway connections. Our BE scanning also identified underappreciated functional hotspots, including an allosteric site in the KRAS N-terminus. We also identify catalytically impaired CRAF mutants that confer bidirectional resistance and sensitization to structurally similar MEK inhibitors, distinguishing functional differences between structurally related compounds. These findings establish BE scanning as a scalable approach for network-level chemical-genetic interaction mapping, revealing new mechanistic insights into one of the most studied oncogenic pathways.

## Introduction

Advances in functional genomic methods have enabled massive, unbiased studies of protein variants in living cells. Among these, base editor (BE) scanning offers the versatility and scalability of CRISPR-based screening but with amino acid-level resolution to evaluate variant effect in a cellular environment. Base editors typically consist of a Cas9 nickase fused to an adenine or cytosine deaminase (ABE or CBE respectively), which catalyze deamination of DNA bases to generate transition mutations within an editing window specified by programmable sgRNAs^1,2^. Using sgRNA libraries tiling genes of interest, BE scanning can test thousands of mutations in a pooled format. This approach has been deployed on diverse proteins to interrogate their function, including BRCA1/2^3^, DNMT3A^4,5^, the spliceosome complex^6^, IFNγ pathway genes^7^, cyclin-dependent kinases^8^, and assorted oncology targets such as BRAF, KRAS, and PARP^9^. These BE scans have identified drug resistance mutations, clarified variant effects, and uncovered mechanisms of protein regulation. Yet, given the advantages of BE scanning, such as scalability, endogenous editing, and minimal cellular engineering requirements, its potential to dissect more complex biological systems remains largely untapped.

BE scanning presents a compelling approach to investigate cell signaling pathways where multiple genes, including redundant or non-redundant paralogs, operate within coordinated networks. One such pathway is the RAS-RAF-MEK-ERK axis or MAPK signaling cascade, which regulates cell division and is frequently activated in human malignancies^10^. MAPK signaling has been the subject of intense study and is the target of numerous small molecule therapeutics. Inhibitors of key pathway members, including KRAS, BRAF, and MEK1/2, are FDA approved, and many other compounds targeting the pathway are currently in clinical development^11–13^. Although efficacious in many contexts, inhibitors of MAPK signaling face significant challenges, including the emergence of drug resistance often due to mutations that reactivate signaling in cells^14^. The recurrence of similar resistance mechanisms across drug classes suggests that resistance may reflect properties of pathway organization, rather than liabilities of individual targets or compounds alone^15,16^. An additional challenge is the complex mechanistic features underlying many MAPK inhibitors which remain not fully characterized. For example, mutant-selective BRAF inhibitors can paradoxically activate wild-type RAF, and several MEK inhibitors have variable potency depending on the presence or absence of RAF and KSR^17,18^. With these challenges in mind, we set out to leverage BE scanning with chemical inhibitor profiling to systematically investigate the MAPK pathway, with the goals of dissecting drug mechanisms of action (MOA) and resistance on a pathway-wide level — laying the groundwork for potentially generalizable strategies to interrogate other cell signaling pathways.

Here we performed large-scale BE scanning of 22 MAPK-signaling pathway genes. To interrogate the functional effects of MAPK mutations in various contexts, we applied multiple selection pressures including dropout comparing early and late timepoints, pathway hyperactivation, pathway inhibition at various signaling nodes, and specific inhibition of MEK with diverse compounds. Together, our results comprise a chemical-genetic map of MAPK signaling, revealing how drug resistance and sensitization signatures differ depending on both the node of inhibition and drug MOA. The full dataset is available in an interactive format at https://liaulab.github.io/2026_MAPK/. Strikingly, we find that most resistance-conferring base edits target hotspots distal to the inhibitor binding pocket and often in proteins other than the target protein altogether. These resistance hotspots frequently coincide with somatic mutation hotspots observed in treatment-naïve cancer patients, consistent with broader pathway activation being a common drug-resistance mechanism. Leveraging these resistance signatures, we show that chemical-genetic profiles can recover aspects of known pathway organization, providing a framework to identify key pathway connections and relationships between proteins. At a granular level, BE scanning also revealed novel regulatory sites and resistance mechanisms within individual pathway members including a previously unrecognized KRAS N-terminus site that regulates sensitivity to pathway inhibition. Lastly, we uncover a class of CRAF mutations that confer bidirectional resistance/sensitivity to MEK-targeting compounds, revealing functional differences in compound MOA and pointing to CRAF as a key determinant of MEK inhibitor potency. Together, our study provides a pathway-level analysis of drug action and resistance across MAPK signaling and establishes BE scanning as a powerful approach for systems-level chemical-genetic analysis and functional site discovery across a cell signaling pathway.

## Results

### Base editor scanning of the MAPK pathway

We used CRISPR BE scanning to mutagenize thousands of amino acids across the MAPK pathway and assess their relative fitness in diverse contexts. To this end, we designed a large-scale BE library to tile 22 core and accessory pathway genes with adenine and cytosine base editors (ABE8e and BE3.9, respectively) (**Fig. 1a,b**). To achieve greater mutational depth, we used base editors incorporating SpG Cas9, which recognizes an expanded NGN PAM. The resulting library includes 15,406 sgRNAs predicted to make 12,888 unique missense mutations across 8,183 amino acid positions (combining predicted edits from CBE and ABE). We also included additional control sgRNAs: 200 non-targeting sgRNAs and 100 sgRNAs predicted to mutate splice sites of essential genes. For each screen, we transduced ABE or CBE sgRNA libraries into H358 (KRAS^G12C^) cells, applied a selection pressure, and then quantified changes in sgRNA abundance. Z-scores for each sgRNA were calculated where positive scores indicate higher sgRNA abundance compared to a control arm and negative scores indicate lower sgRNA abundance (see Methods). As expected, sgRNAs targeting splice-sites of essential genes were depleted (negative Z-score) relative to non-targeting sgRNAs, confirming the activity of CBE and ABE in our BE scans (Extended Data Fig. 1d).

**Fig. 1:**
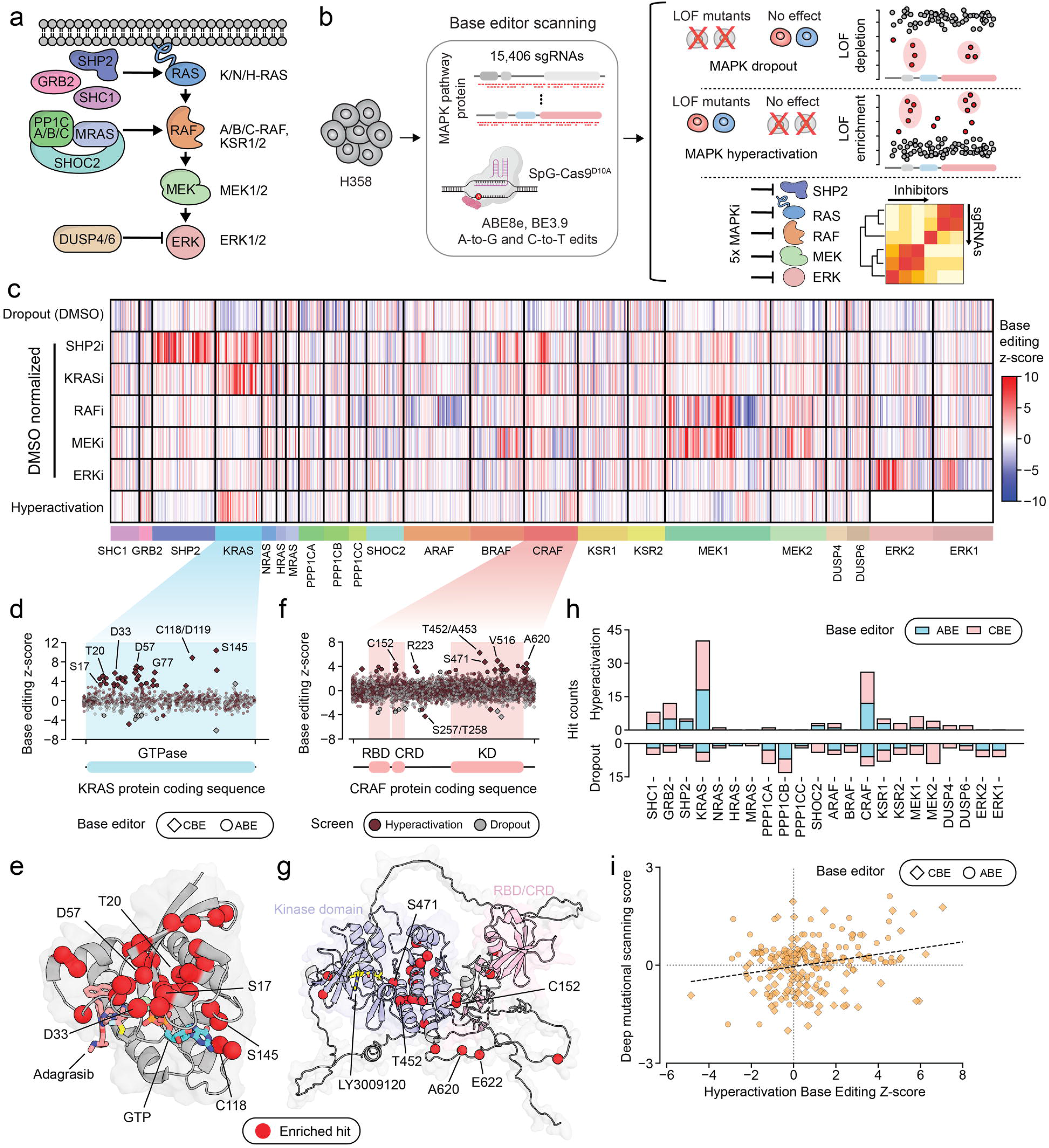
Pathway-wide base editor scans of the MAPK signaling cascade. **a**, Schematic of the MAPK signaling cascade and the 22 genes included in our pathway-wide BE scanning library. **b**, Overview schematic of BE dropout scan, LOF hyperactivation scan, and five-node inhibitor resistance scan. **c**, Heatmap of BE scanning scores schematized in **b** showing all sgRNAs that are hits (|Z-score| > 3) in one or more treatment conditions. The sgRNAs are grouped by gene and ordered within each gene according to that sgRNA’s position on the protein coding sequence. Negative Z-scores indicate loss-of-function in dropout mode, while positive hits indicate loss-of-function in hyperactivation mode. For the five pathway inhibitors, higher Z-scores indicate enrichment compared to DMSO control corresponding to inhibitor resistance. **d**, Scatter plot showing Z-scores of sgRNAs targeting the KRAS protein coding sequence from ABE and CBE hyperactivation positive-selection and dropout negative-selection loss-of-function scans in H358 cells. Negative Z-scores indicate loss-of-function in dropout mode, while positive Z-scores indicate loss-of-function in hyperactivation mode. Scores were calculated by comparing sgRNA abundance after 15 days between doxycycline-treated and untreated groups or untreated and an early time point sample for hyperactivation and dropout respectively. Solid dots indicate sgRNAs with |Z-score| > 3. Select sgRNAs are labeled with the amino acid they are predicted to mutate. **e**, Structure of KRAS (PDB: 6UT0) with red spheres indicating residues predicted to be edited by sgRNAs enriched in hyperactivation-mode BE scan (Z-score > 3). **f**, Scatter plot as in **d** showing Z-scores of sgRNAs targeting the CRAF protein coding sequence. CRAF RAS binding domain (RBD), cysteine rich domain (CRD), and kinase domain (KD) are labeled. **g**, Structure of CRAF (AF-P04049-F1-model_v6) with active site inhibitor, LY3009120, aligned from a structure of BRAF (PDB 5C9C). Residues are shown in spheres that are predicted to be edited by sgRNAs enriched in hyperactivation-mode BE scanning (Z-score > 3). **h**, Stacked bar graph showing hit sgRNA counts identified in hyperactivation scan (Z-score > 3) and dropout scan (Z-score < -3) for all screened genes shown on the positive and negative y axes, respectively. **i**, Scatter plot showing correlation between KRAS BE hyperactivation scan scores and independent KRAS deep mutational scanning data^22^ (Pearson 0.3042, p value < 0.001) Each point in the plot is an sgRNA. A DMS score is calculated for each sgRNA by taking the average of all the DMS Z-scores for the predicted mutations of a given sgRNA.

### Loss-of-function screening identifies functionally essential sites

To identify loss-of-function (LOF) sgRNAs targeting the MAPK pathway we used two selection modes, dropout and hyperactivation. Dropout screening identifies sgRNAs that install mutations which impair essential processes and are thus depleted from the population over time. Dropout scores are obtained by comparing untreated cells grown for 15 days to the same population of cells on day 0. For hyperactivation screening, we took advantage of the sensitivity of H358 cells to activation lethality whereby overexpression of active KRAS dramatically impairs proliferation^19^. This phenomenon enables an enrichment-based screen (i.e., up-assay) for LOF base edits, as LOF mutations enable cell survival in this context by decreasing pathway signaling. We reasoned that an up-assay would be more sensitive than dropout screening, because it allows even a small population of successfully edited cells to proliferate and dominate over time. Conversely, the signal observed in dropout screens is directly limited by incomplete editing, so only LOF sgRNAs with high editing efficiency are identified as hits. For this purpose, we used engineered H358-tetO-KRAS^G12V^ cells, whose growth is arrested following doxycycline-induced KRAS^G12V^ expression unless MAPK signaling is inhibited either chemically or genetically^19^. Hyperactivation BE scanning was conducted by introducing BE libraries into H358-tetO-KRAS^G12V^ cells and then growing these cells for 15 days in the presence or absence of doxycycline. Hyperactivation BE scanning Z-scores for each sgRNA were calculated by comparing the doxycycline treated and untreated groups such that a positive Z-score is assigned to LOF sgRNAs. We performed a pilot screen tiling ERK2 and identified numerous enriched sgRNA hits in ERK2 that correlate with LOF mutations previously identified by deep mutational scanning (**Extended Data Fig. 1b,c**). Because ERK2 was previously identified as a top hit in a genome wide knockout screen in this cell line, we were concerned that in a pooled library spanning the full MAPK pathway, ERK2-targeting sgRNAs might dominate and mask weaker signal elsewhere. We therefore excluded sgRNAs targeting ERK1/2 in the full-scale hyperactivation screen.

Our hyperactivation-based BE scanning identified numerous sgRNA hits (Z-score > 3), especially in core pathway members such as KRAS and CRAF (**Fig. 1d,f**). We observed similarities in the regions identified in both dropout and hyperactivation modes such as residues 56-59 and 145 in KRAS. sgRNAs targeting these residues were enriched in the hyperactivation mode and depleted in dropout mode, indicating that these sgRNAs install mutations that disrupt KRAS function (**Fig. 1d**). Many of the strongest hyperactivation-mode hits identified in KRAS target essential motifs identified in prior literature. For example, sgKRAS-S17 is predicted to edit the phosphate-binding P-loop, and a similar mutation, S17N, has been shown to arrest NIH 3T3 cell proliferation by trapping RAS in a GDP-bound state (**Fig. 1d,e**)^20^. Additionally, an sgRNA targeting KRAS D33 is strongly enriched in our hyperactivation screen. D33 falls within the switch I effector loop and is essential for high-affinity effector binding; nearby mutations Y32R or T35A substantially decrease affinity for the RAS-binding domain of RAF^21^. We also observed a statistically significant correlation between KRAS sgRNA scores and an independent deep mutational scan that identified mutations that suppress oncogenic KRAS activity providing orthogonal support that these sgRNAs introduce LOF mutations^22^ (**Fig. 1i**). In CRAF, we found that hyperactivation-enriched sgRNAs map to residues with defined structural or regulatory roles, that we would expect to be essential for protein function. For instance, sgCRAF-C152 is predicted to mutate a structural, zinc-coordinating cysteine in the cysteine rich domain, a region that is required for RAS engagement^23^ (**Fig. 1f,g**). Moreover, several hit sgRNAs, including sgCRAF-A620 and sgCRAF-E622, are predicted to edit near pS621, a phosphorylation site that is required for CRAF folding, stability, and activation^24^.

Hyperactivation screening identified many hit sgRNAs in KRAS and CRAF which are genes known to be essential in KRAS-driven cancers^25,26^. The greater number of hits identified in these two genes by hyperactivation suggests a greater sensitivity of enrichment-based selection in this experimental setting (**Fig. 1h**). However, we also found that dropout scanning identified more hits dispersed across the pathway, potentially reflecting non-MAPK related functions that may preclude sgRNAs from being enriched in our hyperactivation scan. For example, in PPP1CA/B, LOF sgRNAs were more prevalent in the dropout mode but rare in the hyperactivation mode. PPP1C plays a role in several essential cellular processes beyond regulation of RAF which would explain why LOF mutations in PPP1C may impair cell growth in dropout screens and fail to enrich during MAPK-specific hyperactivation^27^. Altogether, our hyperactivation screening approach provides a complementary and, in some cases, more sensitive method for identifying base edits that impair protein function.

### Base editing identifies drug-resistance and sensitizing mutations

We next sought to chart chemical-genetic interactions throughout the pathway using the same BE-scanning libraries but expanded to include ERK1/2-tiling sgRNAs. For chemical perturbations, we selected five well-characterized inhibitors targeting five distinct nodes of the pathway: batoprotafib (SHP2i), adagrasib (KRASi), LY3009120 (RAFi), trametinib (MEKi), and temuterkib (ERKi) (**Fig. 1b**). For comparability, we dosed each inhibitor at a moderate cellular potency (∼EC_60_-EC_90_) as determined in 5-day growth assays (**Extended Data Fig. 1a**). We introduced base editor libraries into H358 cells and then exposed them to selection pressure (inhibitor) or vehicle control (DMSO) for two weeks before assessing sgRNA abundances by next generation sequencing and calculating sgRNA Z-scores.

These BE scans, spanning inhibition at five different pathway nodes, yielded numerous sgRNA hits scoring as drug-resistant (Z-score > 3) or drug-sensitizing (Z-score < -3) across genes and inhibitor conditions (**Fig. 1c**). Due to the large number of sgRNA hits identified, we employed a pooled approach for initial validation. We designed a validation library consisting of enriched and depleted hit sgRNAs (|Z-score|>3), non-editing controls, and splice-site/stop-codon sgRNAs for each of the 22 MAPK pathway genes. Using this library, we conducted a similar screen in H358 cells using the same five inhibitors at the previously used concentrations but extended to 18 days of compound treatment (**Fig. 2a**, see Methods). Validation screen results strongly correlated with our initial BE scans (**Fig. 2b**). Notably, sgRNAs conferring resistance were more reproducible than those conferring sensitization, indicating the greater robustness of enrichment-based selection.

**Fig. 2:**
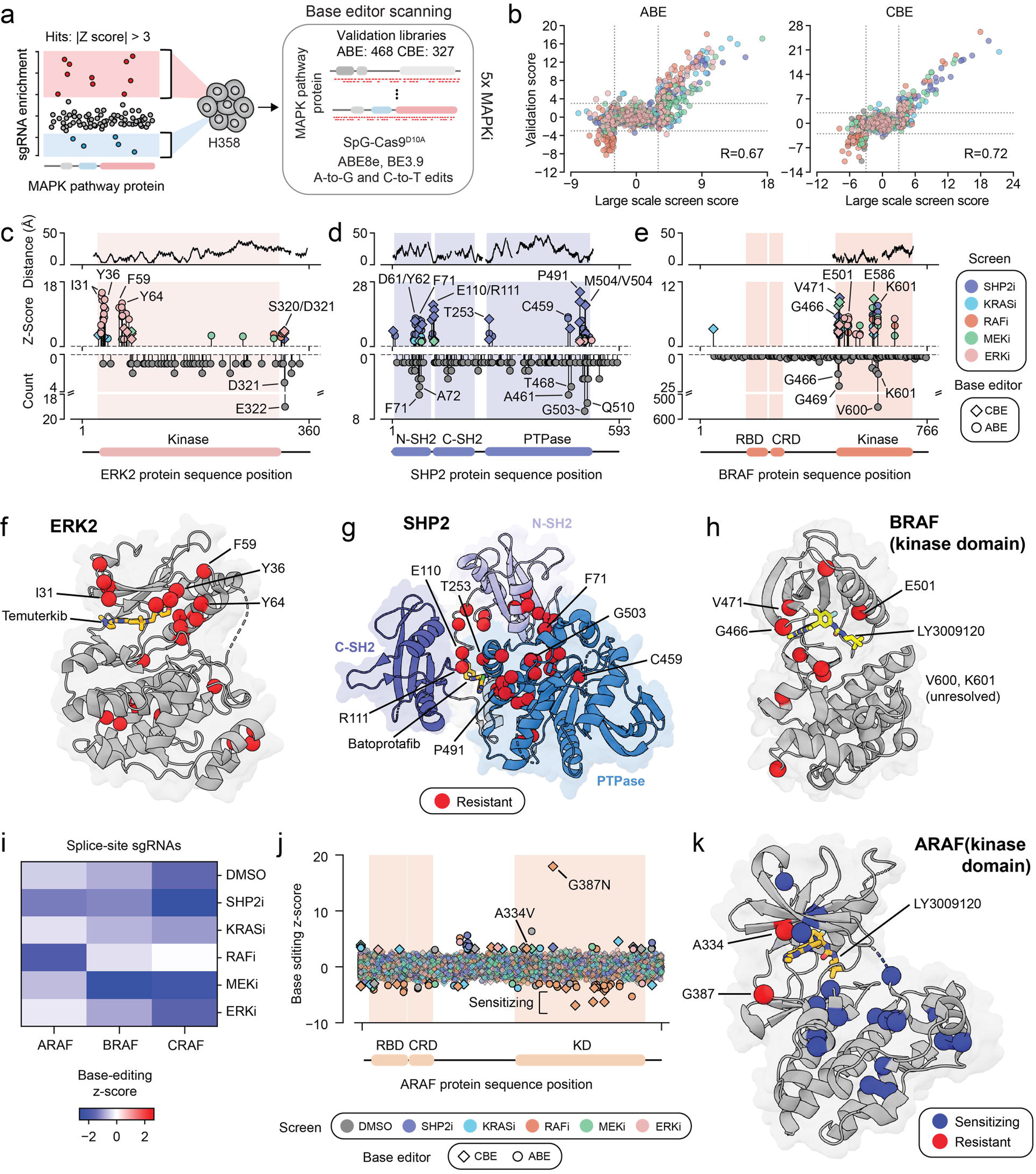
Inhibitor-resistance BE scanning reproducibly identifies functional sites that align with known biology. **a**, Schematic of BE scanning five-node inhibitor validation screen. **b**, Scatter plots comparing large-scale, inhibitor-resistance BE scans and validation screens for both ABE (Pearson R=0.67) and CBE (Pearson R=0.72). Each point in the plot is an sgRNA. Large-scale screen scores are Z-scores, and validation scores are control-referenced Z-scores calculated using the mean and standard deviation of the non-targeting controls. **c-e,** Plots for ERK2 **c**, SHP2 **d**, and BRAF **e** showing sgRNAs with Z-scores > 2 in the validation screen (middle y-axis) along each protein coding sequence. Each axis also shows somatic mutation counts by amino acid position from cBioPortal (bottom y-axis). Distance of each residue from the corresponding inhibitor derived from the PDB structures shown in f-h (top y-axis). **f-h,** Residues predicted to be edited by validated resistance hits shown as red spheres on respective structures. **f,** ERK2 (PDB: 6RQ4) **g**, SHP2 (PDB: 7JVM) **h**, BRAF (PDB: 5C9C). **i,** RAF-targeting splice-site and premature-stop-codon sgRNA mean Z-scores for each condition normalized to day 0 (n=5 for ABE, n=5 for CBE, all combined). RAFi synergizes specifically with ARAF in contrast to BRAF and CRAF. **j**, Scatter plot showing Z-scores of sgRNAs targeting the ARAF protein coding sequence from ABE and CBE large-scale, inhibitor-resistance BE scans. Positive Z-scores indicate inhibitor resistance while negative Z-scores indicate sensitization. Z-scores associated with inhibitors were calculated by comparing sgRNA abundance in an inhibitor treated group to the DMSO treated group. DMSO scores were calculated by comparing the DMSO treated group to the day 0 population. Solid dots indicate sgRNAs with |Z-scores| > 3. Two resistance sgRNAs validated in the follow-up validation screen, sgARAF-A334 and sgARAF-G387. **k,** Structure of ARAF (PDB: 9AXM) with LY3009120 aligned from an inhibitor-bound BRAF structure (PDB: 5C9C). Residues predicted to be edited by validated sgRNAs are shown as spheres.

To understand how sgRNA hits may confer drug resistance, we measured, for each hit sgRNA, the distance between each hit sgRNA’s predicted edited residue(s) and the corresponding inhibitor’s binding site within the target protein (**Fig. 2c–e and Extended Data Fig. 2a,b**). This analysis revealed that some sgRNA hits are predicted to edit residues close to the inhibitor binding site — likely leading to mutations that disrupt compound binding. For example, many sgRNA hits in ERK2 that confer resistance to ERKi target residues near the inhibitor binding site (**Fig. 2c (top axis), 2f**). sgRNAs targeting I31, Y36, F59, and Y64 were strongly and specifically enriched in the ERKi condition — all of these residues are proximal to the temuterkib (ERKi) binding site, and have previously been identified as hotspots for ERKi resistance by deep mutational scanning^28^. Likewise, sgRNAs targeting SHP2 at E110/R111, T253, and P491 conferred strong specific resistance to SHP2i (batoprotafib) and mutate residues close to the compound (**Fig. 2d,g**). These examples are consistent with the notion that mutating residues proximal to the inhibitor binding site is a common mechanism of drug resistance to targeted therapies.

However, a large proportion of resistance sgRNA hits did not target residues proximal to the inhibitor binding sites but instead sites distal (**Fig. 2d,e**). Many of these distal site-targeting sgRNAs overlap with clinical mutation hotspots, suggesting that mutations that upregulate signaling often provide drug resistance (**Fig. 2c–e** and **Extended Data Fig. 2a,c**). For example, SHP2 sgRNAs editing at or near the clinical hotspots, F71 or G503, are strongly enriched in cells treated with SHP2i and to a lesser extent by other pathway inhibitors (**Fig. 2d,g**). Interestingly, we also identified two strong resistance sgRNAs predicted to mutate the SHP2 active site cysteine 459 to arginine, likely impairing SHP2’s catalytic activity. While it seems counterintuitive that gain-of-function (GOF) SHP2 variants could be catalytically inactive, emerging evidence suggests that the scaffolding function of SHP2 can supersede its phosphatase activity in some contexts^29^. Prior studies of pathogenic SHP2 variants also reveal discrepancies between SHP2 catalytic activity and its cellular functions^30,31^. Our results parallel these findings and suggest that SHP2 phosphatase activity is dispensable in H358 cells treated with SHP2i.

We also found overlap between clinical mutation hotspots and sgRNAs that were strongly resistant to pathway inhibition in BRAF. BRAF G466, G469, and K601 are mutational hotspots in human cancer, and we found sgRNAs which target near these sites can promote resistance to various pathway inhibitors (**Fig. 2e,h**). Clinically observed mutations at these sites are known to disrupt autoinhibitory interactions in the BRAF kinase domain and facilitate signaling by promoting active RAF dimers^32,33^. This suggests that broader pathway activation, rather than direct disruption of compound binding, can contribute to resistance.

Additionally, our BE scans identified resistance and sensitizing sgRNAs targeting ARAF that provide insight into the action of RAFi in cells. We observed that sgRNAs predicted to knock out ARAF, as well as sgRNAs targeting the ARAF kinase domain synergize with the RAFi, LY3009120, yielding strong negative Z-scores (**Fig. 2i,j and Extended Data Fig. 2b**). Synergy has also been reported for similar RAF inhibitors when paired with ARAF knockdown.^25^ Together, this suggests that cells depend more on ARAF when CRAF and BRAF are inhibited, and by extension that LY3009120 comparatively spares ARAF despite initial reports of pan-paralog potency^34–36^. Interestingly, our validation screen confirmed two ARAF-targeting sgRNAs that specifically confer resistance to RAFi. These sgRNAs are predicted to make A334V and G387N mutations respectively, both of which are proximal to the modeled LY3009120 binding site (**Fig. 2j,k and Extended Data Fig. 2c**). Previous work also identified mutations of ARAF G387 in cell culture models and patients that were strongly resistant to another RAFi, Belvarafenib^37^. These in-target resistance mutations indicate that LY3009120 likely engages ARAF at the concentration used in our BE scan. Taken together, these data are consistent with a partial-inhibition model, whereby RAF inhibitors impede cell growth to the extent that they inhibit ARAF, which can otherwise compensate for the inhibition of CRAF and BRAF. Moreover, our findings demonstrate how BE scanning can recover known functional sites and resistance/synergy mechanisms, giving us confidence that our approach could also reveal novel biology.

### Mutations near the KRAS N-terminus regulate its activity

Next, we asked whether our drug-resistance-BE-scanning revealed novel functional sites in core pathway members. Among the validated resistance sgRNAs targeting KRAS we found an unusual hit in the N-terminus of KRAS, sgKRAS-Y4, which is predicted to install the compound mutation Y4C/K5G/E (**Fig. 3a**). This mutation is distal from both the adagrasib and GTP-binding pockets and confers resistance to both SHP2i and KRASi (**Fig. 3b,c**). Moreover, a similar, *KRAS^K5N^* mutation has been observed in a small number of RASopathy patients and is expected to be GOF, motivating us to investigate the effects of sgKRAS-Y4^38^. By introducing this sgRNA individually in H358 cells, we confirmed its editing activity and found that it generates two compound mutations, Y4C+K5G and Y4C+K5E (Extended Data Fig. 3b). We also confirmed the ability of sgKRAS-Y4 to confer resistance to KRASi and SHP2i in a competitive growth assay (**Fig. 3d and Extended Data Fig. 3a**).

**Figure 3:**
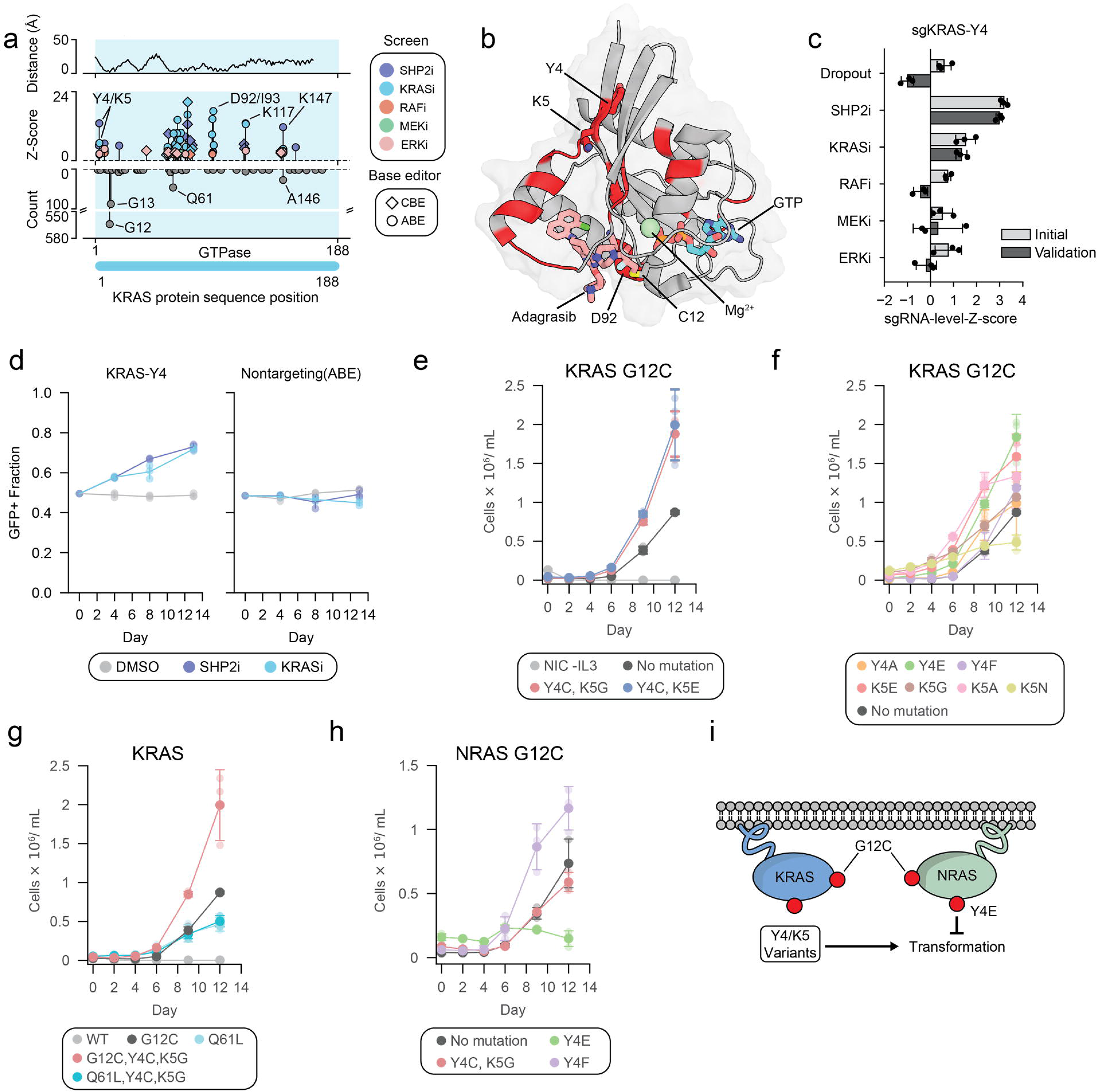
Residues Y4 and K5 regulate activity and inhibitor sensitivity in KRAS G12C. **a,** Plot for KRAS showing hit sgRNA Z-scores above Z-score > 2 in the validation screen (middle y-axis), somatic mutation counts from cBioPortal (bottom y-axis), and distance in Å of each residue from adagrasib in an inhibitor-bound KRAS structure (PDB: 6UT0) (top y-axis). **b,** Validated resistance hits represented in red on the adagrasib-bound KRAS structure (PDB: 6UT0). Residues 4 and 5 are distant from both ligand-binding pockets. **c,** Bar plot of individual sgRNA Z-scores (not averaged across replicates) for sgKRAS-Y4 in the initial BE scan and validation screen. **d,** Single sgRNA competitive growth assay for sgKRAS-Y4 in the presence of DMSO, adagrasib (30nM), or batoprotafib (3μM). Data are presented as mean +/- standard deviation (n = 3). **e,** Ba/F3 transformation assay measuring cell count over time following IL3 withdrawal with KRAS variants in *cis* with G12C expressed from a strong promoter (EF1α). Data are presented as mean +/- standard deviation. (n = 3) **f,** Ba/F3 transformation assay as in **e** with single mutations of KRAS in cis with G12C. **g,** Ba/F3 transformation assay as in **e** testing Q61L alone or combined with Y4C+K5G in *cis*. **h,** Ba/F3 transformation assay as in **e** with NRAS variants expressed from a strong promoter (EF1α). **i,** Cartoon schematic summarizing the results of Ba/F3 transformation assays. KRAS mutants at residues Y4 and K5 increase transformation in combination with G12C. The phospho-mimetic mutation, Y4E, has opposite effects in NRAS and KRAS.

Because H358 cells edited with sgKRAS-Y4 are resistant to upstream SHP2 inhibition, we hypothesized that these mutations stimulate signaling rather than modulate inhibitor binding. However, it was unclear whether Y4C/K5G/E functioned in *cis* or *trans* with the G12C allele, which is heterozygous in H358 cells. Furthermore, the relative contribution(s) of the Y4C, K5G, and K5E mutations to the resistance phenotype were also unclear. To disentangle these effects, we assessed the transforming potential of various KRAS N-terminus mutations alone and in combination with G12C when introduced to Ba/F3 cells. In this assay, proliferation in the absence of IL3 is a surrogate for RAS activity and MAPK pathway activation. When expressed from a weak or strong promoter, KRAS^G12C^ transforming potential was increased by Y4C+K5G and Y4C+K5E mutations (**Fig. 3e, Extended Data Fig. 3d**). However, these N-terminus mutations had no effect in KRAS*^WT^* (**Extended Data Fig. 3c,d**). Moreover, many other single mutations at position 4 or 5 increased KRAS*^G12C^* activity but also have no effect in a KRAS*^WT^* background (**Fig. 3f, Extended Data Fig. 3f,g**). These findings align with published deep mutational scanning data of KRAS showing that mutations at positions 4 or 5 confer drug resistance or promote transformation but only in *cis* with G12C^15,22,39,40^ (**Extended Data Fig. 3e**). We also found that K5N did not increase Ba/F3 transformation, either by itself or in combination with G12C (**Fig. 3f, Extended Data Fig. 3f,g**). Interestingly, Y4C/K5G had no additional effect when combined in *cis* with another oncogenic mutation, Q61L which is known to ablate KRAS GTPase activity (**Fig. 3g**). This shows that the effect of Y4/K5 substitutions depends on the specific biochemical state created by G12 RAS mutations.

Previous literature indicates that RAS proteins can be negatively regulated via phosphorylation at Y4, which directs mono/di-ubiquitination of RAS by RABGEF1^41–43^. To our knowledge, this mode of regulation has only been observed for NRAS, HRAS and Drosophila RAS, not KRAS. We found that Y4 of KRAS can regulate signaling but does so uniquely. The phosphomimic mutation Y4E in KRAS strongly promotes transformation while the phospho-null mutation, Y4F, has a more mild effect (**Fig. 3f, Extended Data Fig. 3f**). In contrast, the same mutations in NRAS have opposite effects — Y4E attenuates NRAS^G12C^ activity while Y4F augments it (**Fig. 3h, Extended Data Fig. 3h**). The specificity of this epistasis is striking considering that the N-terminal residues 1-86 of KRAS and NRAS are identical. Together, these data indicate functional dependency between the N-terminal region of RAS, oncogenic G12 mutations, and the non-conserved residues of KRAS/NRAS (**Fig. 3i**). This example illustrates the value of BE scans as a starting point for discovery — nominating a functional site that can be brought into focus with orthogonal assays.

### High-level analysis reveals patterns in base editing sgRNA resistance

Having established the utility of our BE scanning to uncover individual functional sites, we next sought to investigate resistance across all pathway members at a systems level. High-level structural analysis revealed that while some validated resistance sgRNAs target residues proximal to the inhibitor binding site or elsewhere in the respective drug target (i.e., in-target), most residues targeted by resistant sgRNAs were found in other pathway members entirely (**Fig. 4a**). We refer to the latter as “out-of-target” resistance sgRNAs. Based on a simple chemical epistasis framework, we expected that most resistance mutations would occur at or downstream of an inhibited node, because upstream activating mutations would be masked by the action of downstream inhibitors. This pattern held true for KRASi and RAFi and unavoidably for SHP2i because we did not include upstream RTKs (**Fig. 4b and Extended Data Fig. 4b and Supplementary Table 2**). Interestingly, MEKi and ERKi were susceptible to resistance sgRNAs targeting upstream pathway members. We speculate that this could be due to moderate inhibitor dosing, which may tune down signaling, but allow strong upstream activation to overcome inhibitor action.

**Fig. 4:**
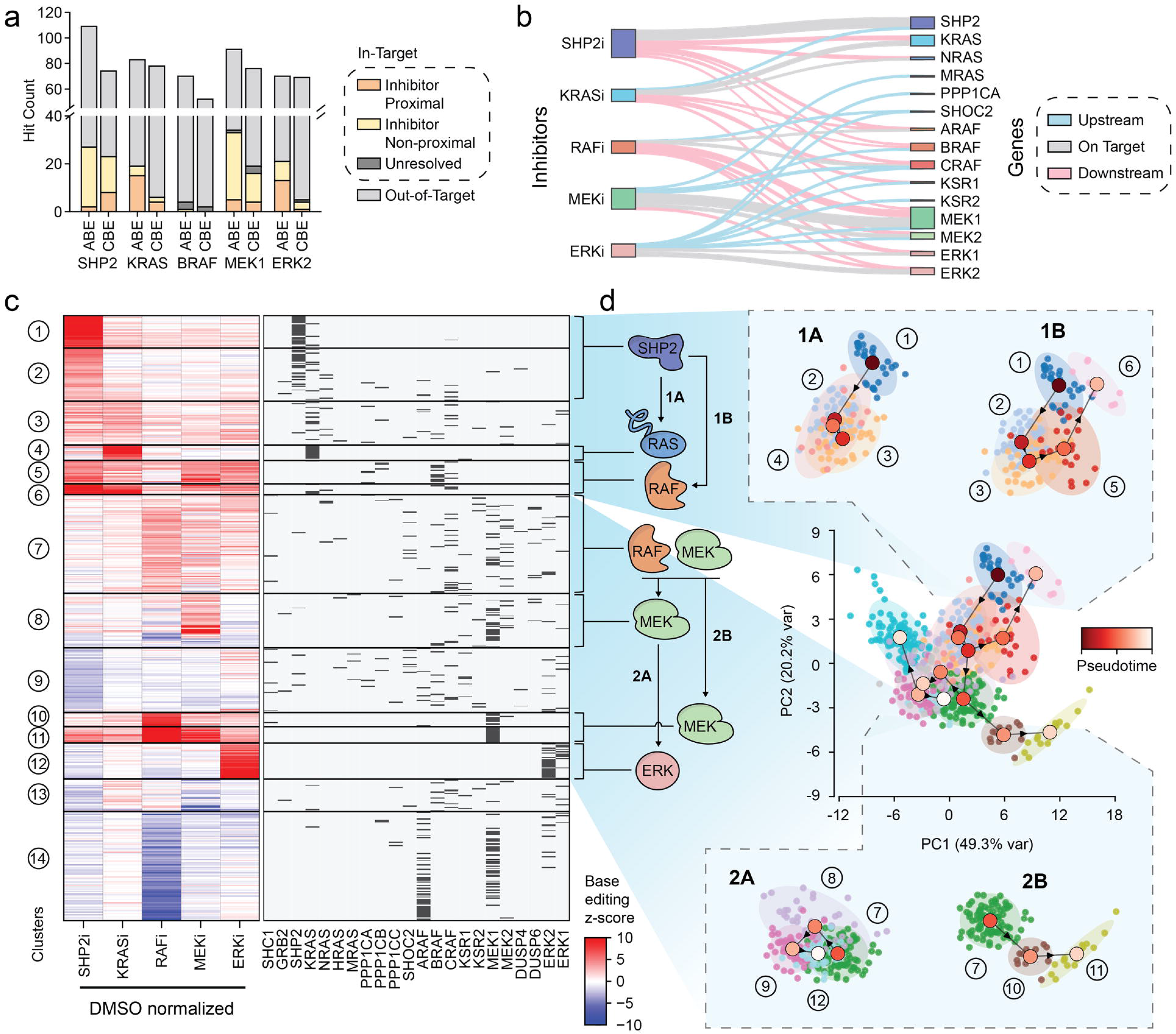
Pathway-wide analysis of base editor scanning. **a,** Bar plot summarizing locations of sites targeted by validated resistance sgRNAs using PDB structures. Inhibitor proximal sgRNAs target amino acids within 4 Å of the respective inhibitor, inhibitor non-proximal sgRNAs edit amino acids in the protein target but >4 Å from the inhibitor, and unresolved sgRNAs target residues in regions not resolved in the referenced PDB structure. The rest of the sgRNA hits shown are resistant to the indicated drug but edit genes other than the direct inhibitor target and are referred to as out-of-target. **b,** Sankey plot showing the number of validated hit sgRNAs (control-referenced validation Z-score > 1) from each inhibitor condition and indicating the protein they edit. Connections with fewer than 2 sgRNAs are not shown. The width of each band indicates the number of validated resistant sgRNAs which appear for that inhibitor and target that protein. A wider band indicates more hit sgRNAs. MAPK proteins downstream of the inhibitor target are shown in red, while proteins upstream are shown in blue. In-target resistance sgRNAs are shown in grey. **c,** GMM clustered heatmap of hit sgRNAs from the validation screen (|control-referenced Z-score| > 2). Clusters are numbered on the left of the heatmap, and genes targeted by each sgRNA are indicated in hashes on the right. The far right is a cartoon where each cluster is represented by the protein(s) that is most targeted by the sgRNAs in that cluster. Arrows between cartoon proteins correspond to the trajectories described in **d**. **d,** All sgRNA shown in the heatmap summarized in PCA space, with a trajectory between clusters defined by Minimum Spanning Tree (MST). Specific sub-trajectories 1A, 1B, 2A, 2B are called out on the top and bottom correspond to specific parts of the PCA which emphasize connections between clusters and their dominant genes.

Because each sgRNA has a unique enrichment signature described in 5 chemical dimensions, we decided to use a clustering approach to group functionally related sgRNAs and evaluate the relationships between them. We chose Gaussian mixture model (GMM) clustering, which probabilistically groups sgRNAs by their five-node inhibitor resistance profiles by assuming the data fits to a set of Gaussians (**Fig. 4c**). Of the resulting clusters, several are characterized by sgRNAs that confer specific resistance to one inhibitor, and these sgRNAs predominantly edit the respective protein targeted by that compound (i.e., in-target). For example, Cluster 12 is defined by highly specific resistance to ERKi, as all 31 sgRNAs in the cluster target ERK1/2. Similarly, inhibitor-specific clusters were identified for SHP2i (Clusters 1 and 2), KRASi (Cluster 4), and MEKi (Cluster 8) and were largely in-target. Mapping these clusters onto protein structures shows that most inhibitor-specific resistance sgRNAs edit residues near an inhibitor binding site (**Extended Data Fig. 4c–e**). These corresponding mutations likely reduce inhibitor affinity to confer resistance.

This analysis also revealed sgRNA clusters that confer broad resistance to several compounds. For example, Cluster 5 primarily comprises BRAF/CRAF-targeting sgRNAs that confer resistance to most inhibitors tested and to a lesser extent the RAFi, while Cluster 10 is composed mostly of MEK1/2-targeting sgRNAs that confer preferential resistance to the RAFi. In-target RAFi resistance in BRAF/CRAF may target residues proximal to the inhibitor binding site while, in contrast, out-of-target RAFi resistance in MEK1/2 may activate the pathway downstream. Similarly, Cluster 6 is populated by KRAS and BRAF/CRAF-editing sgRNAs which confer resistance to the SHP2i and KRASi. Finally, Cluster 11 comprises MEK1/2-targeting sgRNAs that confer resistance to the RAFi and MEKi. These patterns suggest that activating mutations at the node of inhibition or one node downstream are especially common mechanisms to overcome small molecule inhibition in MAPK.

### Pathway-wide analysis recapitulates key connections in the signaling cascade

After characterizing phenotypically similar clusters of sgRNAs, we tested whether chemical-genetic interactions embedded within base-editing resistance profiles could be used to computationally infer key interactions and relative pathway order. Since many resistance sgRNAs are likely to reactivate pathway signaling, we predicted that sgRNAs would exhibit chemical-epistasis relationships with the pathway inhibitors such that resistant sgRNAs would be more enriched by upstream inhibitor treatment rather than downstream inhibitor treatment. Thus, resistance profiles of sgRNAs might encode a chemical-genetic position within the pathway. We adapted TSCAN, a method used to infer pseudo-time with scRNA-seq data, to construct a minimum spanning tree (MST) connecting sgRNA clusters identified by the GMM.^44^ GMM clustering accommodates the noisy nature of sgRNA resistance profiles, similar to scRNA-seq, by calculating a probability for each sgRNA across all clusters to build a continuous trajectory. A “pseudo-pathway” position for each cluster can be inferred by projecting it onto the MST, and the resulting cluster order is used to generate an ordered representation of the resistance profiles. By considering the greatest proportion of sgRNAs within each cluster, we assign some clusters as being dominated by one protein. Then, we asked whether this ordering recapitulated known relationships between key pathway members. Cluster 1, which corresponds to in-target SHP2i inhibition, was chosen as the starting node of the trajectory based on literature evidence that SHP2 regulates KRAS.^45^ This analysis inferred a pathway trajectory, which we broke down into sub-trajectories for interpretability (**Fig. 4d**).

The trajectory between clusters recapitulates several connections in the SHP2-RAS-RAF-MEK-ERK cascade (**Fig. 4d**) with sub-trajectories that identify key interactions such as SHP2-KRAS or RAF-MEK-ERK. Trajectory 1A highlights the transition from SHP2i resistance in SHP2 (Clusters 1 and 2) to KRASi resistance within KRAS (Cluster 4) consistent with the established placement of KRAS downstream of SHP2 and identifying a connection between SHP2 and KRAS. Trajectory 1B highlights a different transition from SHP2i resistance mediated by in-target base edits in SHP2 (Clusters 1 and 2) to a broader pan-resistance phenotype driven by activating mutations in downstream proteins such as KRAS and BRAF/CRAF (Clusters 5 and 6) (**Fig. 4d and Extended Data Fig 4f**). Trajectory 2A starts at Cluster 7 and highlights MEKi-specific resistance in MEK1/2 (Cluster 8) and ERKi specific resistance in ERK1/2 (Cluster 12), placing ERK1/2 downstream of MEK1/2 which are downstream from BRAF/CRAF. Trajectory 2B branches toward stronger RAFi/MEKi resistance driven by sgRNAs targeting MEK1/2 (Clusters 10 and 11), identifying a strong RAF-MEK connection (**Fig. 4d and Extended Data Fig. 4g**). Trajectory 2C includes the cluster with the most sensitizing phenotype where mutations in ARAF/MEK/ERK sensitize cells towards RAFi (**Extended Data Fig. 4h-i**). Notably, trajectory 2C stands out as the most functionally distinct, and its placement near the end of the trajectory underscores a limitation of MST inference: all clusters are assigned a position in the trajectory even when their interactions aren’t clearly interpretable. In summary, trajectories 1A/2A describe the specific in-target resistance profiles while trajectories 1B/2B describe the path from specific resistance to broader pan-resistance within a SHP2–KRAS and RAF–MEK couple. By assuming continuity in chemical-genetic interactions, the resulting ordering recovered several established relationships within the MAPK cascade and type of analysis may have broader applicability to the study of other systems for inferring key connections between pathway members.

### Base editing reveals distinct types of resistance to MEK inhibitors

We next explored whether BE scanning can parse and classify functional differences in the mechanisms of action (MOA) of closely related compounds. Specifically, we investigated MEK inhibitors because they have been reported to differ in their MOAs (**Fig. 5a**). Most MEK inhibitors are allosteric type 3 kinase inhibitors, binding to a site adjacent to the ATP-binding pocket in a noncompetitive fashion. Despite binding the same pocket, allosteric MEK inhibitors diverge in their effects on MEK phosphorylation and MEK–RAF complex formation^46^ (**Fig. 5b**). Trametinib and cobimetinib disrupt MEK–RAF heterodimers, while avutometinib and trametiglue stabilize an inactive MEK–RAF complex.^18,47,48^ Other allosteric MEK inhibitors like selumetinib and mirdametinib also increase MEK–RAF interaction, but simultaneously promote MEK phosphorylation and allow for pathway reactivation.^49^ More recently, several orthosteric MEK inhibitors have been developed, including MAP855, which directly compete with ATP binding.^50^

**Fig. 5:**
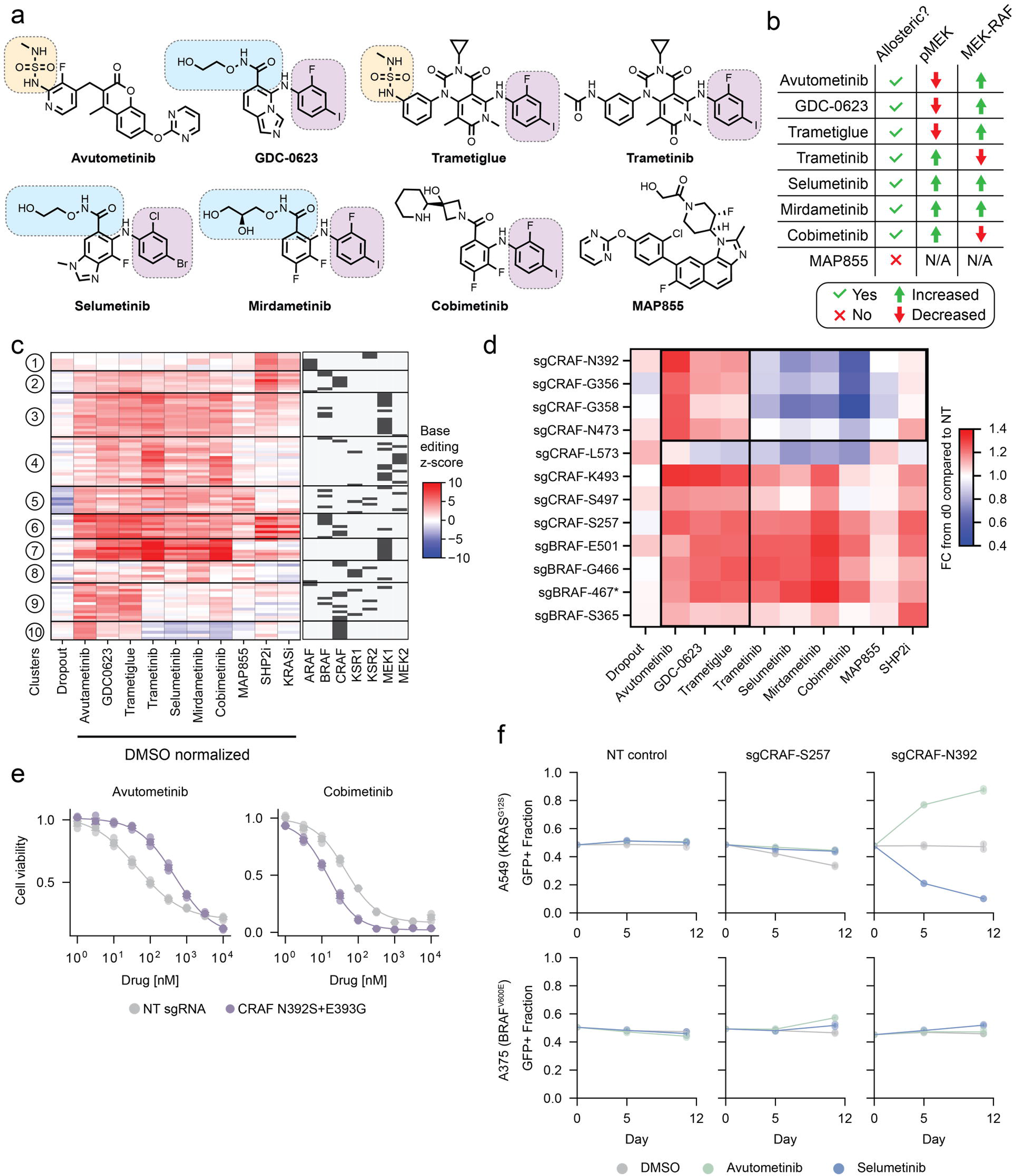
Base editor scanning functionally differentiates structurally similar MEK inhibitors. **a,** Structures of all MEK-targeted compounds used for MEK inhibitor-focused BE scanning. Similar functional groups across compounds are highlighted. **b,** Table summarizing MEK inhibitor traits based on literature evidence. pMEK column indicates the effect of each compound on levels of cellular phosphorylated MEK^48,49,58,62,73^. MEK-RAF column summarizes the effect of each compound on MEK-RAF complexes in cells, as determined by co-IP^18,48,49^. N/A indicates that we were unable to find experimental evidence in the literature. **c,** GMM clustered heatmap of resistance sgRNAs from the MEK inhibitor-focused BE scanning screen (Z-score > 4 in at least one condition) shown by their resistance profiles across all MEKi and for SHP2i and KRASi from previous five-node BE scanning screen. Clusters are numbered on the left of the heatmap, and genes targeted by each sgRNA are indicated in hashes on the right. **d,** Single sgRNA validation (competitive growth assay) for select CRAF and BRAF-targeting sgRNAs performed with a single endpoint on day 12. Heatmap shows mean of 3 replicates normalized to day 0 and then to normalized to a nontargeting control. See methods for inhibitor concentrations. **e,** Four-day growth assay (CellTiter-Glo) with base-edited homozygous clone carrying N392S+E393G mutation or a single cell clone carrying a nontargeting control sgRNA. Performed in triplicate and normalized to DMSO treated wells. Data are presented as mean and standard deviation. (n = 3). **f,** Single sgRNA validation (competitive growth assay) in A549 and A375 cells. Data are presented as mean +/- range (n = 2).

We reasoned that mechanistically distinct MEK inhibitors should give rise to distinct resistance mutation signatures. To that end, we performed BE scanning using a subset of our larger library to tile MEK1/2, ARAF, BRAF, CRAF and KSR1/2 with both CBE and ABE, and eight different MEK inhibitors (**Fig. 5a and Extended Data Fig. 5a**). As before, we dosed each compound at approximately equivalent potencies (∼GI_75_) determined in 5-day growth assays (**Extended Data Fig. 5b**). We then grouped sgRNAs that scored as strongly resistant (Z-score > 4) in at least one MEK inhibitor condition by Gaussian Mixture Model clustering. To help differentiate types of resistance mechanisms, we included the prior screen resistance sgRNA scores for the KRASi and SHP2i conditions in the clustering analysis (filtering Z-score > 4) (**Fig. 5c**). We found that sgRNAs in Clusters 1 and 2 conferred resistance to upstream MAPK inhibitors but not MEK inhibitors, suggesting general pathway reactivation rather than specific perturbation of MEK inhibitor binding. We found these clusters included sgRNAs predicted to introduce known activating mutations such as CRAF S257P which disrupts an inhibitory 14-3-3 binding motif,^51,52^ ARAF S214P which perturbs an analogous motif,^53^ or BRAF E501G, a GOF mutation which has been reported in RASopathy patients.^54^ Cluster 6 sgRNAs confer resistance to both upstream inhibition and all allosteric MEK inhibitors but not the orthosteric inhibitor, MAP855, and were also predicted to introduce known activating mutations. For example, sgBRAF-K601 is predicted to introduce the BRAF K601G mutation, which is similar to oncogenic mutations in BRAF, K601E/N/T.^33^

Our analysis also identified Cluster 7 sgRNAs that confer resistance to allosteric MEK inhibitors with minimal resistance to upstream inhibition. This includes sgMEK1-L115, sgMEK1-H119 and sgMEK1-E203. L115 and H119 are located in the C helix of MEK1 near the allosteric inhibitor binding site^18^. In vitro kinase assays show that the L115P mutation completely ablates potency of another MEK inhibitor, PD184352, indicating that mutations in this helix can disrupt small molecule binding while preserving MEK1 catalytic activity.^55^ The specific resistance conferred by sgMEK1-G202, predicted to install E203K and G202K mutations, is less understood. Prior literature suggests that E203K activates MEK1 rather than disrupting drug binding; however, full interpretation of this sgRNA is complicated by the predicted co-mutation of G202.^56^ Notably, strong in-target resistance sgRNAs were not enriched by MAP855, in agreement with independent results showing that MAP855 retains activity against an array of allosteric MEK inhibitor-resistance mutations.^50^

Unexpectedly, sgRNAs in clusters 9 and 10 selectively conferred resistance to three MEK-targeting compounds, avutometinib, trametiglue, and GDC-0623, but not to other allosteric MEK inhibitors (**Fig. 5c**). Cluster 10 sgRNAs even synergize with other MEK inhibitors. This bidirectional effect of the sgRNAs is striking considering the similar chemical structures and binding poses of many of these compounds.^18,46^ Trametiglue and trametinib differ only by a sulfonamide extension, and GDC-0623 and selumetinib are structurally related and share a similar binding mode (**Fig. 5a**). However, these two groups of MEK inhibitors exhibit sharply bifurcated responses to cluster 9 and 10 sgRNAs, indicating that these MEK inhibitors operate by different mechanisms. Based on prior literature, the closest correlate separating these two groups of inhibitors is the ability to suppress rather than induce pMEK in cells (**Fig. 5b**).

### CRAF kinase domain mutations confer bidirectional resistance and sensitization to MEK inhibitors

To evaluate these bidirectional MEK inhibitor signatures, we individually tested the effects of 20 hit sgRNAs using competitive growth assays in the presence of DMSO, the eight individual MEK inhibitors, or SHP2i (**Fig. 5d and Extended Data Fig. 5c**). This group of sgRNAs included strong hits from clusters 9 and 10 as well as several control sgRNAs as comparators, such as those that confer pan-resistance or pan-sensitization. Of these, 13/20 sgRNAs robustly validated, providing strong resistance or sensitivity by competitive growth. Four bidirectional sgRNAs robustly conferred resistance to three MEK inhibitors while sensitizing to others and all four edit the kinase domain of CRAF (**Fig. 5d and Extended Data Fig. 5c**). Targeted amplicon sequencing confirmed successful editing by these 4 sgRNAs, which introduce N392S/E393G, G356K, G358N, and N473S/I474V mutations (**Extended Data Fig. 5d**). We next generated a clonal population of H358 cells carrying a homozygous mutation of N392S/E393G (**Extended Data Fig. 5e**) and compared to a clonal cell line transduced with a non-targeting sgRNA. *CRAF^N392S/E393G^* cells were resistant to avutometinib and sensitized to cobimetinib across a range of inhibitor doses (**Fig. 5e**). To assess the generalizability of the bidirectional CRAF mutations, we tested sgCRAF-N392 in two other cell lines, A549 and A375 cells. A549 cells, which carry a mutant *KRAS^G12S^*allele, exhibited the same bidirectional phenotype (**Fig. 5f**). In contrast, sgCRAF-N392 did not have a substantial effect in A375 cells, which harbor *BRAF^V600E^*, suggesting that the phenotype depends on a KRAS-mutant cellular context.

Notably, the validated bidirectional CRAF sgRNAs have a unique phenotype compared to that of other sgRNAs predicted to make S257P, K493G, or S497G mutations in CRAF which uniformly confer resistance to all inhibitors tested (**Fig. 5d**). The bidirectional CRAF sgRNAs also do not phenocopy the effects of sgCRAF-L573, which sensitizes to most tested compounds, potentially by installing a LOF or destabilizing mutation into CRAF. Interestingly, an sgRNA predicted to generate BRAF^E501G^, which is analogous to CRAF^E393G^, confers a pan-resistance phenotype rather than a bidirectional MEK inhibitor-specific effect (**Fig. 5d**). This shows that the bidirectional phenotype is distinct from other classes of functional sgRNAs which confer resistance or sensitivity more generally and that it is likely paralog specific.

Alignment of BRAF and CRAF sequences reveals that the four validated, bidirectional CRAF base edits loosely resemble class 3 BRAF mutations, suggesting a mechanistic explanation for their effects (**Fig. 6a**). Class 3 BRAF mutants, including G464V/E, G466V/E/A, and N581S, have impaired catalytic activity but promote signaling in a RAF-dimerization-dependent manner.^33^ By analogy, we hypothesized that bidirectional CRAF mutations may also impair kinase activity while preserving or promoting dimerization. The locations of these substitutions are consistent with this hypothesis (**Fig. 6b**). E393 is a conserved residue that interacts with the catalytic lysine in the active state and with the inhibitory turn of the activation loop in the inactive state; G356 and G358 are in the glycine-rich loop proximal to ATP; and N473 interacts with Mg^2+^ in the catalytic pocket. We therefore predicted that mutation of any of these sites would impair CRAF catalytic activity.

**Fig. 6:**
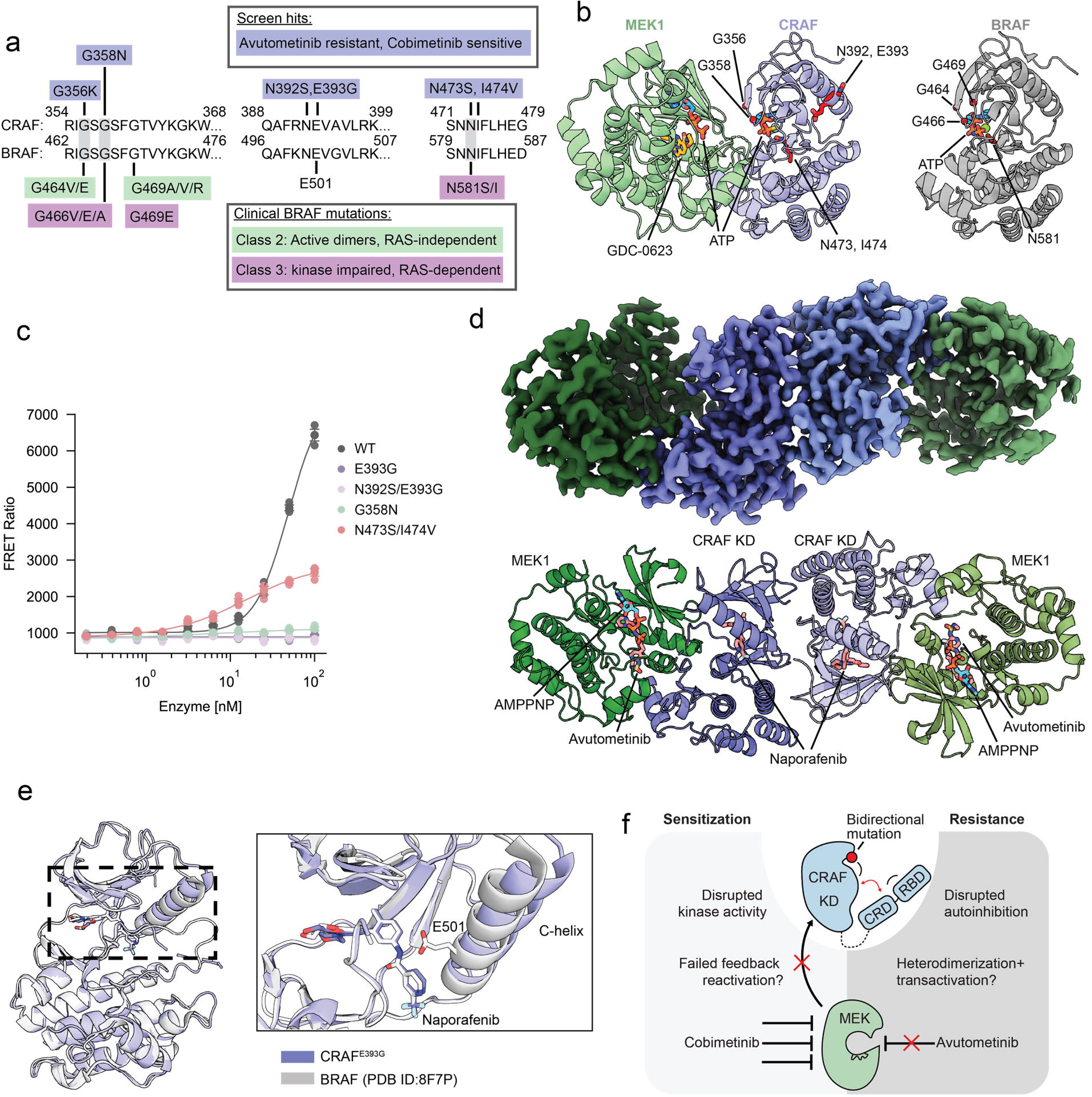
Biochemical and structural investigation of bidirectional CRAF mutations. **a,** Amino acid sequence alignments of BRAF and CRAF indicating class 2/3 clinical BRAF mutations and bidirectional CRAF mutations identified via BE scanning. **b,** Mutation sites from **a** shown in red on a structure of CRAF-MEK (PDB:9MMQ) and BRAF-MEK (PDB:6V2W, only BRAF shown) structures. **c,** TR-FRET-based kinase assays measuring MEK1 phosphorylation by CRAF kinase domains containing the indicated mutations. Data are presented as mean and standard deviation, with replicate values shown (n=3) measuring kinase activity of the indicated mutants. **d,** Cryo-EM structure of the heterotetrameric CRAF^E393G^/MEK1 complex with naporafenib, avutometinib, and AMPPNP. The cryo-EM density map is shown in the upper panel, with the corresponding atomic model displayed as ribbons in the lower panel. CRAF^E393G^ is colored blue and MEK1 is colored green. Naporafenib, avutometinib, and AMPPNP are shown as stick representations. **e,** Structural comparison of the CRAF^E393G^/MEK1 (this study) and BRAF in the complex with naporafenib (PDB ID: 8F7P). **f,** Working model of how bidirectional CRAF mutations affect sensitivity to MEK-targeted compounds. Left: bidirectional mutations impair CRAF kinase activity which prevents reactivation of CRAF following release of negative feedback resulting in greater potency of cobimetinib and similar MEK inhibitors. Right: bidirectional mutations disrupt the autoinhibited conformation of CRAF which may alter heterodimerization to confer resistance to avutometinib and similar MEK inhibitors.

To test this hypothesis, we purified CRAF kinase domains harboring several of these bidirectional mutations and quantified the kinase activity of each against recombinant MEK1. All tested bidirectional CRAF mutations reduced or ablated catalytic activity (**Fig. 6c**). Despite their compromised catalytic activity in vitro, these CRAF mutants did not measurably impair cell growth in our BE scans or competitive growth assays, nor did their corresponding sgRNAs score as LOF in our hyperactivation screen (**Fig. 1f, 5f, and Supplementary Table 4**). These observations align with prior studies showing that A549 and other KRAS-driven lung cancer cells do not depend on CRAF kinase activity^25,26,57^.

A potential explanation for the synergy between bidirectional CRAF mutations and MEK inhibitors is that feedback reactivation of CRAF serves a protective role in cells challenged with MEK inhibitors, and this feedback is ablated when CRAF catalytic activity is impaired. Prior work shows that either CRAF knockdown or a type II RAF inhibitor similarly synergize with selumetinib or mirdametinib in KRAS-mutant cells^49,58^. However, because RAF inhibitors can inhibit all RAF paralogs, it is difficult to ascribe this synergy specifically to CRAF inhibition rather than ARAF or BRAF inhibition. To investigate further, we used a low dose of AZ628, a RAF inhibitor which achieves partial-selectivity toward CRAF (3 nM AZ628 inhibits CRAF > 75%, BRAF <25%, ARAF ∼0%)^36^. In H358 cells, we found that 3 nM AZ628 strongly synergizes with cobimetinib, but up to 10 nM AZ628 does not further inhibit proliferation of cells treated with avutometinib (**Extended Data Fig. 6a**). Importantly, both the N392S/E393G mutation or low dose of AZ628 synergized with cobimetinib, but when combined, they had little to no additional effect on H358 proliferation, consistent with overlapping mechanism (**Extended Data Fig. 6b**). Together, these data suggest that CRAF catalytic activity is largely dispensable during basal growth but contributes to survival when cells are challenged with certain MEK inhibitors.

### Biochemical and structural characterization of mutant CRAF/MEK complexes

The converse phenotype — resistance of bidirectional CRAF mutants to some MEK inhibitors — is not intuitively explained by loss of CRAF catalytic activity. We initially hypothesized that because avutometinib and trametiglue are both RAF-MEK glues, bidirectional CRAF mutations might oppose the action of these compounds by weakening the CRAF-MEK interaction. However, SPR using immobilized mutant CRAF kinase domain did not detect substantial differences in CRAF-MEK affinity, either in the presence or absence of MEK inhibitors (**Extended Data Fig. 6c**). Because SPR did not reveal a major change in MEK-CRAF kinase domain affinity, we next examined the structural features of mutant CRAF.

To understand the possible impact of these mutations on protein structure, we sought to determine cryo-EM structures of CRAF^E393G^/MEK complexes in both the autoinhibited (heterodimeric) and active (heterotetrameric) states using cryogenic electron microscopy (cryo-EM). Co-expression of MEK and CRAF^E393G^ in insect cells yielded a heterodimeric complex as previously observed for CRAF^WT^. We were unable to obtain a single-particle reconstruction of this complex due to conformational heterogeneity, likely due to destabilization of the regulatory C-helix (**Extended Data Fig. 7a**). In addition to its conserved role in catalysis, E393 plays a key role in stabilizing the C-helix out-conformation of the kinase in the inactive state (**Extended Data Fig. 7b**). Thus, we expect that loss of this stabilizing interaction increases the dynamics of the C-helix, leading to the observed particle heterogeneity.

In contrast, we were able to obtain a high-resolution single-particle reconstruction of the “active” MEK:CRAF:CRAF:MEK complex with the E393G variant (**Fig. 6d and Extended Data Fig. 8**). In this preparation, we added the RAF inhibitor, naporafenib, which induces RAF dimerization to form a heterotetrameric complex. The resulting structure revealed a conformation like that previously observed for heterotetrameric CRAF^WT^/MEK complexes^59,60^. In particular, the C-lobe contacts between the MEK and CRAF kinase domains are preserved (**Extended Data Fig. 7c**). Differences in inhibitors bound in the CRAF active site preclude detailed comparison of the N-lobe conformations between these CRAF structures. However, comparison with the naporafenib-bound structure of the active BRAF dimer reveals a closely similar conformation. The most apparent difference is a modestly outwardly displaced C-helix in the CRAF^E393G^ mutant structure, apparently due to loss of a hydrogen bond between E393 (which lies in the C-helix) and the central amide of the inhibitor (**Fig. 6e**). The cryo-EM map is well-resolved in the region of the mutation and bound inhibitor and supports loss of this interaction (**Extended Data Fig. 7d**). Together, these structural data indicate that E393G preserves the core RAF-MEK dimerization interfaces while modestly increasing C-helix dynamics. If these dynamics increase CRAF dimerization with other RAFs, especially ARAF, this might explain the resistance to MEK inhibitors, as recent evidence shows that ARAF-activated-MEK is less sensitive to inhibition than MEK activated by other RAF paralogs^61^. However, the complete mechanistic basis for the inhibitor-specific resistance pattern requires further biochemical dissection to establish.

## Discussion

In this work, we demonstrate how pathway-scale BE scans can be paired with diverse selection pressures to understand chemical-genetic relationships throughout the MAPK cascade. Furthermore, we exploited the multiple chemical dimensions of our dataset to perform pathway-wide analyses including clustering and minimum spanning tree analysis to recapitulate several key connections within the pathway. Ultimately, we hope a similar approach could be extended to map poorly understood pathways or expanded to characterize all putative effectors of MAPK signaling.

BE scanning allowed us to identify novel functional sites and generate hypotheses about their mechanisms by drawing from the multiple chemical dimensions of our dataset. For example, we found that sgRNAs that confer specific resistance to KRASi often target residues in the switch II pocket, likely because these mutations impair KRASi binding (**Extended Data Fig. 4c**). In contrast, sgKRAS-Y4 mutates an allosteric, N-terminal sequence, conferring resistance to both KRASi and SHP2i via general upregulation of KRAS activity (confirmed through orthogonal assays). Interestingly, follow-up experiments revealed an epistatic relationship between mutations at position 4-5, G12C, and the non-conserved residues of RAS paralogs. Further investigation such as characterization of altered binding partners or localization might clarify how this N-terminal regulatory site modulates RAS and whether this mechanism could be exploited pharmacologically.

BE scanning also surfaced several counterintuitive features of MAPK signaling that are only partially explained in the literature. For example, we found that CRAF mutations that ablate kinase activity have no impact on cell proliferation, a phenomenon that has been described several times and likely stems from CRAF’s primary role of dimerization rather than catalysis in KRAS-mutant lung cancer^25,57^. We also found sgRNAs that are resistant to SHP2i but are predicted to edit the catalytic cysteine to arginine, C459R. This aligns with emerging literature showing that SHP2’s scaffolding role in recruiting SOS1 may supersede its enzymatic function in some contexts^29^.

Patterns in sgRNA resistance also provided insight into RAF inhibitor MOA. We found that ARAF knockout synergized with RAFi, while two sgRNAs targeting residues near the ARAF active site provide specific resistance to RAFi, consistent with a partial inhibition model of ARAF by LY3009120. Interestingly, we found many BRAF and CRAF-targeting sgRNAs that provide strong resistance to several pathway inhibitors but generally fail to resist LY3009120 (Cluster 5 **Fig. 4c**). For example, an sgRNA that makes the activating, CRAF S257P mutation, confers resistance to most pathway inhibitors but substantially less resistance to LY3009120. We imagine two possible explanations for this pattern. First is that orthosteric inhibitors may be generally insensitive to many but not all resistance mutations. We observed a similar trend for MAP855, the only orthosteric MEK inhibitor we tested, which was impervious to sgRNAs that conferred resistance to allosteric MEK inhibitors and upstream inhibitors. Another possible explanation is that variants that induce the active, dimerized conformation of RAF are actually more easily inhibited by RAFi, a phenomenon that has been highlighted in recent work^62,63^. We also note that LY3009120 is a RAF dimer inhibitor, and we would likely have obtained a different result with RAF monomer inhibitors such as vemurafenib, in a sensitive context, for which RAF activation/dimerization is a well-known mechanism of resistance^64^.

Finally, we used BE scanning to evaluate differences in MEK inhibitor MOA and found a unique class of mutations in the CRAF kinase domain that differentiate between two groups of MEK inhibitors. These bidirectional variants are catalytically impaired and resemble class 3 BRAF mutations, which disrupt autoinhibition and promote signaling through heterodimerization^32,33^. It is plausible that bidirectional CRAF mutations also disrupt autoinhibition and promote dimerization with ARAF to achieve resistance. However, more study is required to determine the exact mechanism for this resistance and why similar mutations in BRAF, like E501G, behave differently. Importantly, avutometinib, which received FDA approval in 2025, was strongly resisted by these bidirectional CRAF mutations^65^. If analogous mutations emerge clinically in patients receiving avutometinib treatment, our data suggest that use of other MEK inhibitors, especially cobimetinib, could be considered rather than assuming class-wide MEK inhibitor resistance.

While adenine and cytosine base editors can only access a small fraction of all possible missense mutations, our work highlights that this depth can interrogate protein domains and identify novel functional sites. We also show that strong selection pressures, i.e., “up-assays” may partially mitigate the challenge of low base editing efficiency, increasing the effective coverage of BE scans. By using hyperactivation as a selection pressure, we conducted an up-assay for LOF variants and identified a greater number of hits in KRAS and CRAF than we identified by dropout scanning. Other cellular contexts are likely needed to extend this hyperactivation strategy to additional proteins in the pathway like NRAS, HRAS, or BRAF, which appear to be less functional in H358 cells. We also envision potential improvements for GOF screening by using a larger library of compounds and testing a range of dose concentrations.

More broadly, our study shows that BE scanning is not merely a means of cataloging variant effect within a single protein, but is an approach for probing the network-level relationships of a signaling pathway. These profiles can generate hypotheses about where resistance emerges, how pathway nodes relate, and how inhibitors differ in mechanistic action. Although MAPK signaling is particularly well suited to this strategy, the general framework should be adaptable to other pathways with tractable cellular phenotypes and available genetic or chemical modulators. We anticipate that pathway-wide chemical-genetic mapping could provide a general framework for dissecting drug mechanism and anticipating resistance, extending naturally to other signaling axes with potent of small molecule modulators. Combining base editing with diverse selection pressures and readouts offers a scalable route to move from functional observation to mechanistic insight for other vital pathways that drive human disease.

## Methods

### Cell culture

All cell lines were cultured at 37 °C and 5% CO_2_ in RPMI1640 or DMEM media (Thermo Fisher Scientific) supplemented with 100U ml^-1^ penicillin and 100 µg ml^-1^ streptomycin (Life Technologies) and 10% Fetal Bovine Serum (Peak Serum). H358, A375, and Ba/F3 cells were a gift from Dr. Matthew D. Shair. H358-tetO-KRAS-G12V cells were a gift from Dr. Arun Unni and Dr. Harold Varmus. A549 (ATCC) cells were a gift from Dr. Andrew Myers. HEK293T cells were a gift from Dr. Brad E. Bernstein. All cell lines were regularly tested for mycoplasma. Ba/F3 cells were supplemented with murine IL3 (PeproTech) prior to transformation assays.

### Plasmid construction

Single sgRNAs were synthesized as oligonucleotides (Genewiz/Azenta), phosphorylated, annealed, ligated into all-in-one base editor vectors which had been previously digested with Esp3I. pRDA_478 (SpG-BE3.9, Addgene, 179096) and pRDA_479 (SpG-ABE8e, Addgene, 179099) were gifts from J. Doench and D. Root. RAS expression plasmids for Ba/F3 experiments were generated via Gibson Assembly using NEBuilder HiFi (New England Biolabs). The weak promoter (hPGK) vector was derived from Cilantro2 (Addgene 74450, a gift form B. Ebert) digested with Esp3I and BamHI to remove GFP. The strong promoter (EF1α) vector was constructed in-house as minimal lentiviral plasmid containing EF1α-RAS-IRES-PuroR positioned between two LTRs. It was digested or PCR amplified prior to assembly with RAS coding sequences. KRAS4b and NRAS coding sequences were PCR amplified from Addgene 159539 and 116767 respectively (gifts from Dominic Esposito or Gordon Mills and Kenneth Scott). All final plasmids were checked via whole plasmid sequencing. (Quintara Biosciences)

### Base editor library design and cloning

The full 22-member library was designed and annotated using the BE-design tool from Ruth Hanna and John Doench (https://github.com/mhegde/base-editor-design-tool), using the ENSEMBL IDs in Supplementary Table 6. BE-hive was used to annotate each sgRNA for predicted editing efficiency. (https://github.com/maxwshen/be_predict_efficiency). Off target scores were calculated using Flashfry (https://github.com/mckennalab/FlashFry). sgRNAs predicted to mutate start codons, the final stop codon, introns, UTRs, splice sites, and those predicted to generate premature stop codons were removed. We also filtered out a small number of duplicate sgRNAs arising from paralog similarity, and a custom script was used to assign the editing site of each sgRNA which was used for plotting enrichment across the length of each gene. 200 nontargeting controls and 100 essential gene splice-donor-targeting controls were included from prior base editing study.^3^ The initial proof-of-concept ERK2 library was designed using CRISPOR (https://github.com/maximilianh/crisporWebsite/) and annotated by adapting existing code (https://github.com/liaulab/DNMT3A_base_editor_scanning/tree/main/library_annotation)

The validation sub-pool library was designed by filtering hit sgRNAs identified in our large-scale, five-inhibitor BE-scan using the following criteria |Z-score| > 3, day 0 sgRNA count > 100, and Hsu2013 specificity score > 20, not intronic. We also selected five sgRNAs for each MAPK gene that edit splice donors, splice acceptors, or that generate premature stop codons. We prioritized sgRNAs with high BEhive scores and those that target splice donors which generally give rise to the most efficient gene knockout.^66^ These validation and knockout pools were designed and cloned separately for ABE and CBE.

sgRNA oligo libraries were barcoded and synthesized as previously described.^67^ (Twist Biosciences). Similar genes were barcoded together to allow focused scans of only select pathway members. Libraries were amplified in two steps. LSPCR1 to amplify a specific barcoded subpool and LSPCR2 to prepare the sub libraries with appropriate Gibson homology arms. Libraries were then incorporated into ABE/CBE vectors that had been digested by Esp3I (ThermoFisher) via Gibson assembly. Assembled DNA was then alcohol precipitated and electroporated into Endura electro-competent cells (Bioresearch Technologies) achieving coverage of > 50x. Colonies were subsequently harvested and plasmid DNA extracted via maxiprep (Qiagen).

### Lentivirus production and transduction

For large scale virus production, 6-8 million HEK293T cells were plated in a 15 cm dish and allowed to adhere overnight. Cells were then transfected using lipofectamine 3000 (ThermoFisher) according to the manufacturer’s protocol with second generation lentivirus packaging plasmids. 9.44 µg of BE plasmid library, 9.44 µg pCMV-VSV-G (Addgene, 8454, a gift from D. R. Liu and P. Salmon) and 18.88 µg pBS-CMV-gagpol (Addgene, 35614, a gift from B. Weinberg) were used. Media was changed 6-8 hours after transfection and virus was collected and filtered through a 0.45 µM filter 36-48 hours later. For single sgRNA validations, validation libraries and KRAS/NRAS ORF expression, virus was generated similarly but with third generation lentiviral system where Gag/Pol genes are split into two separate plasmids, pMDLg/pRRE and pRSV-Rev (Addgene, 12251 and Addgene, 12253, both gifts from Didier Trono). All viral transductions were performed in the presence of 8 μg/ml of polybrene (Santa Cruz Biotechnology) overnight followed by media exchange. Puromycin (Thermo Fisher Scientific) was used at a concentration of 1 μg/ml starting two days after the start of transduction.

### Base editor scanning

Five-inhibitor base editor scans were performed by transducing H358 cells with ABE or CBE library virus at an MOI <0.35 and 500-1000x coverage. Cells were then selected in puromycin for 7-9 days. Following puromycin selection, a day zero pellet was harvested and cells were split into six different treatment groups (DMSO or MAPK pathway inhibitors) in triplicate (18 groups total) where each group had >500x coverage. DMSO treated cells were passaged every 2-4 days. Inhibitor-treated cells had their media changed once on day 7 and split on day 10 or later if they became confluent. On day 14 all cells were harvested.

Validation BE scans were performed similarly but cells were infected at an MOI <0.35 at >1000x coverage, puromycin selection proceeded for 10 days, inhibitor treatments were started at >1000x coverage in triplicate and allowed to proceed for 18 days prior to harvest.

Hyperactivation BE scans were performed similarly but using H358-tetO-KRAS-G12V. Cells were infected at 900-1000x coverage, puromycin selection proceeded for 7 days, and then cells were split into two treatment groups (+/- 1 μg/ml of doxycycline) in triplicate (6 groups total) where each group had 1000x coverage. During doxycycline treatment, cells were continuously exposed to 250 μg/ml of G418 to guard against silencing of the tetO-KRAS-G12V construct. Doxycycline treatment was continued for 15 days prior to harvest.

MEK inhibitor focused BE scans were performed similarly to five-inhibitor base editor scans. Cells were infected at an MOI<0.35 at >1000x coverage and puromycin selection proceeded for 9 days. Cells were then split into 9 different treatment groups (DMSO or MEK inhibitors) in triplicate where each replicate had 500x coverage. DMSO treated cells were passaged every 2-4 days. Inhibitor-treated cells had their media changed once on day 7 and all cells were harvested after 14 days of treatment.

Proof of concept ERK2 scan was also performed in H358-tetO-KRAS-G12V cells and were transduced at an MOI <0.3 and 1000x coverage and puromycin selection proceeded for 6 days. Cells were then split into two treatment groups (+/- 1 μg/ml of doxycycline) in triplicate (6 groups total). Media was refreshed at least every 3-4 days. Cells were harvested after 14 days of Dox treatment.

For all BE scans, gDNA was extracted from cell pellets using either mini or midi-scale QIAamp kits (Qiagen). sgRNA sequences were amplified by PCR from gDNA at 200x coverage or greater (assuming 9.9 pg DNA per cell) using barcoded primers using as previously described (see “sgRNA/shRNA/ORF PCR for Illumina Sequencing”, Broad Institute GPP, https://portals.broadinstitute.org/gpp/public/resources/protocols). PCR products were purified by gel extraction, then pooled and sequenced on Illumina NextSeq or MiSeq systems.

### Base editor scanning analysis

#### Read processing and sgRNA scoring

Fastq files were processed to generate counts of every sgRNA in each screen condition and replicate. Counts for each sample were then normalized and log transformed according to the following equation: log2((count for sgRNA n / total sample sgRNA counts)+1). Replicates were then averaged, and comparisons were made by subtracting log2 normalized counts of one group from another as indicated in the text (e.g. inhibitor x – DMSO).

Read counts and Z-scores for every screen are summarized in Supplementary Table 1, which carries one worksheet per screen and base editor: the positive-selection resistance (GOF) screens, the hyperactivation loss-of-function (LOF) screens, the ERK-focused loss-of-function screen, the MEK inhibitor-focused screens, and the validation screens, each for ABE and CBE. Z-scores were calculated, using the mean and standard deviation of all sgRNAs within each library and base editor. Only the validation screens were normalized using the mean and standard deviation of nontargeting sgRNAs within each library and base editor to generate a control-referenced Z-score.

Annotations were resolved per editor: sgRNAs from ABE screens were assigned the amino acid position, mutation type and list of amino acid substitutions derived from A-to-G editing, and sgRNAs from CBE screens the equivalent values derived from C-to-T editing. sgRNAs whose editing window produces no change to the coding sequence carry a position of −1, as described above, and were excluded from every analysis with an amino acid position axis. Analyses that pool the two editors do so only after this per-editor assignment.

Hit thresholds differ by purpose and are stated with each analysis. Unless noted otherwise, a hit in the primary resistance screens is an sgRNA with |Z| > 3 in the relevant inhibitor-versus-DMSO condition, a validated hit is one with |Z| > 2 in the same condition of the validation screen, a hyperactivation hit in the loss-of-function screens is an sgRNA with Z > 3, and a dropout loss-of-function hit is an sgRNA with a Z < -3.

#### Visualization of BE scanning data

BE scanning data were plotted as sgRNA Z-score against the assigned amino acid position, one panel per protein. sgRNAs annotated as UTR, intronic or splice are excluded from these panels. Domain boundaries for all proteins were manually curated from UniProt or Interpro.

#### Comparison of large-scale BE scanning and validation screens

sgRNAs scoring in the primary resistance screens were re-tested in a focused validation library. For each inhibitor condition and each editor, sgRNAs were called as hits in the primary screen (|Z| > 3) and the sgRNAs in the validation screen that scored (|Z| > 2 in the same condition) were considered validated.

#### Sankey plots of validated resistance hits

To summarize where each inhibitor’s resistance hits fall within the cascade, hit sgRNAs were counted for every inhibitor and protein pair using the validation screen. An sgRNA was counted when its validation control-referenced Z-score was at least 1 in the same condition; ABE and CBE validated hits were summed. Connections between proteins and inhibitors supported by fewer than two sgRNAs were not drawn. Diagrams were drawn with Plotly (v6.9.0) with node positions fixed in MAPK pathway order.

#### Validation screen knockout sgRNA heatmaps

Control-referenced Z-scores for the essential-splice-site or premature-stop sgRNAs targeting MAPK proteins were averaged (ABE n=5, CBE n=5) and these values displayed as a heatmap. For this analysis, all treatment groups were compared to day 0.

#### Structural analysis of in/out of pocket resistance hits

Two sets of structures were used. The first is a set of co-crystal structures of each inhibited node with a representative inhibitor bound: PTPN11 with batoprotafib (PDB 7JVM), KRAS with adagrasib (6UT0), BRAF with LY3009120 (5C9C), MAP2K1 with trametinib (7JUR) and MAPK1 with temuterkib (6RQ4). The second is a set of AlphaFold models (v6 fetched on Oct 1 2025), one for each of the twelve inhibited nodes and their paralogs, with the corresponding inhibitor aligned into the pocket, so that paralogs lacking a co-crystal structure receive their own distance profile. Structures were parsed with the mmCIF parser of Biopython (v1.88) and values are provided in Supplementary Table 5. For every residue of the protein chain, the distance to the ligand was defined as the minimum Euclidean distance between any non-hydrogen atom of that residue and any non-hydrogen atom of the ligand. Residues within 4.0 Å of the ligand were defined as in-pocket residues and further residues were defined as out-of-pocket residues. Hits were taken as sgRNAs with Z > 3 in the condition corresponding to that structure’s inhibitor for both ABE and CBE, and they were classified by their assigned amino acid position. A hit was called in-pocket when an sgRNA targets +/- 1 residues within the defined pocket, out-of-pocket when the sgRNA targets residues outside of the pocket, and unresolved when the sgRNA targets residues unresolved in the structure. The one-residue buffer accommodates the ambiguity in editing site, as base-editors frequently have out-of-window activity.

#### Gaussian mixture model clustering and minimum spanning tree trajectories

sgRNAs were clustered on their 5-inhibitor response profiles with a Gaussian mixture model. sgRNAs with 100 or fewer reads in the Day 0 sample were removed, and non-targeting and essential splice site control sgRNAs were removed so that clusters reflect the MAPK relevant library alone. Z-scores were clipped to the range −10 to 10 in order to reduce the influence of a small number of extreme values on the covariance estimates of the GMMs, and ABE and CBE sgRNAs were pooled, giving 523 sgRNAs described by a five-dimensional response profile. A Gaussian mixture model with full covariance matrices was fitted to these profiles using scikit-learn (v1.9.0), with 14 clusters, five random initializations, and a fixed random seed. A trajectory through the fitted clusters was constructed by joining the cluster means with a minimum spanning tree over their pairwise Euclidean distances (SciPy v1.17.1). One component was designated the root based on known MAPK biology with SHP2 being the start of the MAPK pathway, and each cluster was assigned a pseudotime equal to its cumulative tree distance from the root, obtained by breadth-first traversal and rescaled to the interval 0 to 1. Principal component analysis was used to project the resistance profiles into two dimensions for display only. Clustered heatmaps show sgRNAs as rows, blocked by cluster in trajectory order and ordered by pseudotime within each block.

The MEK inhibitor-focused screen was clustered by the same GMM method but not analyzed as a trajectory. sgRNAs were retained if they showed (Z-score > 4) in at least one of the eight MEK inhibitor conditions or the SHP2i or KRASi inhibitor conditions. A higher cutoff was used to focus our analysis on the strongest hits in the absence of a secondary validation screen. In total 100 sgRNAs were described by an eleven-dimensional resistance profile which was fitted with a ten-cluster Gaussian mixture model.

#### Comparison of LOF sgRNAs to deep mutational scanning data

Base editor scanning results were compared to published deep mutational scanning (DMS) measurements for the same protein, using two datasets: KRAS (HCC827 G12D screen)^22^ and ERK2 (ETP versus DOX)^28^ provided in Supplementary Table 4. DMS datasets were Z-scored prior to comparison. The amino acid substitutions annotated for each sgRNA were matched to the DMS dataset. sgRNAs not predicted to install missense mutations were excluded. Because all three DMS datasets score depletion negatively, an sgRNA making several substitutions was summarized by the average score across its matched substitutions.

#### Comparison to clinical variants

Somatic mutation data for each protein were obtained from cBioPortal (v6 fetched on Aug 27 2025) provided in Supplementary Table 3. The transcript identifiers used are recorded with the analysis code. Missense, nonsense, frameshift insertion, frameshift deletion, in-frame insertion and in-frame deletion mutations were retained, and the residue position of each was parsed from the protein change annotation. Mutations were then grouped and counted per residue.

#### Software and code availability

Analysis was performed in Python (v3.11) using be_scan (v3.0.0) (https://github.com/liaulab/be-scan), NumPy (v2.4.6), pandas (v3.0.5), SciPy (v1.17.1), scikit-learn (v1.9.0), Matplotlib (v3.11.1), Seaborn (v0.13.2), Biopython (v1.88) and Plotly (v6.9.0). All analysis is contained in a single notebook that installs its dependencies and uses input from Supplementary Table 1 to generate the described analyses and plots. The notebook, together with the supporting Python modules, are available at [https://github.com/liaulab/2026_MAPK].

### Cell titer glo proliferation assays

H358 cells were plated in 384 well plates (1,000 cells/well) and treated with inhibitors at a final DMSO concentration of 0.1% and volume of 50 μl. After the indicated number of days, viability was read out by removing 25 μl of media and adding 25 μl of Cell-titer-glo reagent (Promega). Luminescence was quantified on a SpectraMax i3x plate reader.

### Competitive growth assays

H358, A549, and A375 cells were first stably transduced with a GFP expressing lentivirus (Addgene 115643) and bulk sorted to generate +GFP cell lines. GFP expressing cells were transduced with pRDA 478/479 carrying individual sgRNAs of interest or a nontargeting sgRNA. GFP negative cells were transduced with nontargeting sgRNA. All cells were puromycin selected before mixing +GFP cells carrying an sgRNA of interest or nontargeting control with - GFP cells carrying a nontargeting control at a ∼1:1 ratio. Cell mixtures were plated in triplicate wells and treated with inhibitor or DMSO control (0.1% final DMSO concentration). The following concentrations for each inhibitor were used: 1 nM Trametiglue, 10 nM Trametinib, 100 nM-GDC-0623, 300 nM avutometinib, cobimetinib, mirdametinib, 1 nM MAP855, 3 μM Selumetinib, Batoprotafib. Several of these are lower concentrations than used for the BE scan (**Extended Data Fig. 5b**) to ensure sufficient cells were present for flow cytometry on day 12. For MEK inhibitor BE scan validations, cells were measured by a flow cytometry on day 0 of drug treatment and day 12. All flow cytometry data was collected on a NovoCyte 3000RYB flow cytometer (Agilent) and GFP+/- cell counts were assessed on NovoExpress software (v.1.5.0, Agilent). For heatmap summaries, the fold change of each well was calculated between day 12 and day 0. These values were averaged across three replicates and then divided by the mean fold change of the non-targeting control wells. Column labeled dropout in heatmap indicates DMSO treated group. Data for these assays can be for in Supplementary Table 7.

### Single sgRNA genotyping

gDNA was extracted from cells transduced with single sgRNAs or in the case of CRAF-targeting sgRNAs, from cell mixtures harvested on day 12 of competitive growth assay. gDNA was extracted using quick extract solution (Biosearch Technologies) and locus of interest was amplified by PCR using locus-specific primers (see Supplementary Table 8). Barcodes were added via a second PCR before gel extraction and sequencing on lllumina Miseq or NextSeq systems. The resulting fastqs were analyzed using CRISPResso2 (https://github.com/pinellolab/crispresso2) to calculate allelic frequency tables.

### Ba/F3 assays

Ba/F3 cells were transduced with lentivirus and then puromycin selected for 2 days in the presence of IL3. Cells were then washed 3x in PBS to completely remove IL3 and resuspended in RPMI+FBS without IL3. Cells were plated at 1-5 x 10^5^ cells/ml. Cell counts were assessed by flow cytometry on day zero and every 2-5 days thereafter in the presence of helix NP NIR dye to gate out dead cells (Biolegend). Data for these assays can be for in Supplementary Table 7.

### Protein expression and purification

#### CRAF^KD,E393G^/MEK1 complex for the cryo-EM study

Recombinant baculovirus expressing the CRAF kinase domain (residues 337–615) with the E393G mutation, as well as the Y340D/Y341D mutations, was prepared using the pAc8 vector, which contains an N-terminal His_6_-MBP-tag followed by a Tobacco Etch Virus (TEV) protease cleavage site. Recombinant baculovirus expressing the full-length MEK1 (residues 1–393) with the S218A/S222A mutation was prepared using the pAc8 vector, which contains an N-terminal His_6_-tag followed by TEV protease cleavage site. Sf9 cells in liter-scale cultures were co-infected with 0.5% (v/v) of each baculovirus and harvested 3 days post-infection by centrifugation at 1,500g before flash-freezing in liquid nitrogen.

Protein purification was performed as previously described^36^. Briefly, the cells were lysed by sonication in buffer A (50 mM Tris pH 7.5, 300 mM NaCl, 0.5 mM TCEP, 2 mM MgCl_2_, and 50 µM ATP) supplemented with 40 mM imidazole and protease inhibitor cocktail (Thermo Fisher Scientific). Following ultracentrifugation at 40,000 rpm, the clarified lysate was loaded onto a HisTrap HP column (Cytiva). Bound proteins were eluted with buffer A supplemented with 200 mM imidazole and incubated with TEV protease at a 1:50 molar ratio overnight at 4 °C. The sample was subsequently buffer-exchanged into buffer B (50 mM Tris-HCl pH 7.0, 50 mM NaCl, 0.5 mM TCEP, 2 mM MgCl_2_, and 50 µM ATP) and loaded onto a HiTrap SP column (Cytiva) for cation-exchange chromatography. The cleaved His_6_-MBP tag and excess MEK1 were removed in the flow-through, and the remaining proteins were eluted using a linear gradient of buffer A. The eluted proteins were incubated with 200 µM naporafenib, avutometinib, and AMPPNP overnight at 4 °C and then injected onto a Superdex200 increase 10/300 column (Cytiva) equilibrated with buffer containing 20 mM HEPES pH 7.5, 150 mM NaCl, 1 mM TCEP, 5 mM MgCl_2_, 20 µM AMPPNP, 10 µM naporafenib, 10 µM avutometinib. The fractions containing the heterotetrameric CRAF/MEK complex were pooled and concentrated to 1.0 mg/ml using an Amicon Ultra concentrator (30 kDa MWCO, Millipore). The heterodimeric CRAF^E393G^/MEK1 complex in the absence of RAF inhibitor was purified similarly, but without naporafenib.

#### RAF^KD^ proteins for the biochemical studies

Recombinant baculoviruses expressing the BRAF^WT^ kinase domain (residues 423–726) and the CRAF kinase domain (residues 314–618; Y340D/Y341D), including WT, G356K, G358N, E393G, and N473S/I474V variants, were prepared using the pAc8 vector containing an N-terminal His_6_-tag-MBP-tag-TEV cleavage site. Sf9 cells were infected with 1% baculovirus in liter-scale cultures and harvested 3 days post-infection. Cell pellets were lysed by sonication in buffer A (50 mM HEPES pH 7.5, 300 mM NaCl, 1 mM TCEP, 5 mM MgCl_2_, 5% glycerol) supplemented with 10 µM ATPγS, 40 mM imidazole, and protease inhibitor cocktail (Thermo Fisher Scientific). Following ultracentrifugation at 40,000 rpm, the clarified lysates were loaded onto a HisTrap HP column (Cytiva). Bound proteins were eluted using a linear gradient of buffer B (50 mM HEPES pH 7.5, 300 mM NaCl, 1 mM TCEP, 5 mM MgCl_2_, 5% glycerol, 500 mM imidazole). The eluted proteins were injected onto a Superdex200 increase 10/300 column (Cytiva) equilibrated with buffer A. The purified His_6_-MBP-tagged proteins were concentrated to 1–3 mg/ml and used for SPR analysis.

For kinase activity assays, the His-MBP-tag was cleaved by incubation with TEV protease at a 1:50 molar ratio overnight at 4 °C. The cleaved His_6_ MBP-tag and His_6_-tagged TEV protease were removed by passage over a HisTrap HP column (Cytiva). The flow-through containing the tag-free proteins, in buffer A containing 40 mM imidazole, was collected and concentrated to 0.5 – 1 mg/ml for kinase activity assays.

#### MEK1 proteins for SPR analysis

Baculovirus expressing the full-length MEK1 (residues 1–393) with the S218A/S222A mutation was used to infect Sf9 cells. Harvested cells were lysed by sonication in buffer A (50 mM Tris pH 7.5, 150 mM NaCl, 1 mM TCEP, 5 mM MgCl_2_) supplemented with 10 μM AMPPNP, 25 mM imidazole, and protease inhibitor cocktail (Thermo Fisher Scientific). Following ultracentrifugation at 40,000 rpm, the clarified lysates were loaded onto a HisTrap HP column (Cytiva). Bound proteins were eluted using a linear gradient of buffer B (50 mM Tris pH 7.5, 150 mM NaCl, 1 mM TCEP, 5 mM MgCl_2_, 500 mM Imidazole). The eluted proteins were injected onto a Superdex 200 pg 16/600 column (Cytiva) equilibrated with buffer A. The purified proteins were concentrated to 30 mg/ml and flash-frozen in liquid nitrogen.

### Cryo-EM grid preparation and data acquisition

The heterotetrameric CRAF^E393G^/MEK1 complex at a concentration of 0.4 mg/ml was applied to holey gold grids (UltrAuFoil R 1.2/1.3, 300 mesh), which had been glow-discharged for 2 min at 20 mA using GloQube Plus (Quorum). Grids were blotted for 1–3 s using LEICA EM GP (Leica Microsystems) at 11 °C and 95% relative humidity and then immediately plunged into liquid ethane. Grids were initially screened on a Tundra (Thermo Fisher Scientific) operating at 100 keV and subsequently transferred to a Titan Krios (Thermo Fisher Scientific) operating at 300 keV equipped with a Falcon4i direct electron detector and Selectris Energy Filter for data collection. A total of 9,185 micrographs were collected. Movies were recorded with 84 frames per movie at a magnification of 165,000x (corresponding to 0.74 Å per pixel), with a total exposure dose of approximately 50 electrons/Å2 and a defocus range of 0.8 to 1.8 μm. Details of data collection and dataset parameters are summarized in Supplementary Table 9.

### Cryo-EM data processing and model building

All Cryo-EM data processing was performed using cryoSPARC version 5.0.6^68^. Movies were processed using Patch motion correction and Patch CTF estimation. Initial particle picking with a blob size of 80 – 180 Å yielded 3,122,001 particles. Following iterative 2D classification, 387,878 particles were selected for ab initio reconstruction with three classes. Heterogenous refinement followed by homogenous and non-uniform (NU) refinement yielded a map at ∼2.9 Å resolution from 159,747 particles. Using this map, a template was generated and used to pick particles, resulting in 4,487,508 particles. Following iterative 2D classification, 841,229 particles were selected for ab initio reconstruction with three classes. Subsequent homogenous and NU refinement yielded a map at 2.6 Å resolution from 362,796 particles. These particles were further cleaned through multiple rounds of heterogenous refinement with a decoy model. The remaining particles were refined using CTF refinement and reference-based motion correction. A final round of NU refinement yielded a map at 2.53 Å resolution from 305,954 particles. The final map was sharpened using DeepEMhancer. To improve the density of MEK1, local refinement was performed using a mask encompassing MEK1. This locally refined map enabled improved model building of MEK1, which was deposited separately. The overall data processing scheme is summarized in **Extended Data Fig. 8**. For model building, PDB entry 9MMQ was used as an initial model. The model was fitted and subsequently refined using ChimeraX^69^ and Coot^70^, followed by refinement in PHENIX^71^. Final refinement statistics from PHENIX are presented in Supplementary Table 9.

### RAF kinase activity assay

A time-resolved FRET (TR-FRET) assay was conducted to measure kinase activity of RAF proteins using a modified HTRF KinEASE assay kit (Cisbio), as previously described^72^. Briefly, RAF proteins were prepared at concentrations ranging from 100 nM to approximately 50 pM through 2-fold serial dilutions in kinase buffer supplemented with 25 mM MgCl_2_ and 0.25 mM TCEP, along with 250 µM biotinylated-MEK1 as the substrate. Reactions were initiated by the addition of 200 µM ATP. After incubation for 40 min at room temperature, the reactions were quenched with detection buffer supplemented with an anti-phospho MEK1/2 antibody (Revvity) coupled to Eu^3+^ as the FRET donor and XL665-streptavidin conjugate as the FRET acceptor. Following a 30-minute development, FRET signal ratios were measured at 665 and 620 nm using a PHERAstar microplate reader (BMG Labtech).

### Surface plasmon resonance (SPR)

The binding affinities of full-length MEK1 (S218A/S222A) for RAF kinase domain proteins in the presence of various MEK inhibitors (MEKi) were determined by surface plasmon resonance (SPR) using a Biacore 8K instrument (Cytiva). His_6_-MBP-tagged RAF kinase domain variants, including CRAF WT, G356K, G358N, E393G, N473S/I474V, and BRAF WT, were immobilized on a CM5 sensor chip by amine coupling using 0.1 M N-hydroxysuccinimide (NHS) and 0.4 M 1-ethyl-3-(3-dimethylaminopropyl) carbodiimide hydrochloride (EDC) according to the manufacturer’s instructions (Cytiva). For immobilization, RAF proteins were diluted to 20–30 μg/mL in 10 mM HEPES-NaOH (pH 7.5) from stock solutions (1–2 mg/mL). Following immobilization, remaining activated carboxyl groups on the sensor chip were blocked with 1 M ethanolamine (pH 8.5). Reference flow cells were prepared using the same procedure without protein immobilization and were used for background subtraction.

For kinetic analysis, MEK1 was prepared in running buffer containing 10 mM HEPES (pH 7.5), 150 mM NaCl, 1 mM TCEP, 5 mM MgCl_2_, 5 μM AMPPNP, and 0.005% P20 surfactant. For experiments performed in the presence of MEKi, the running buffer was supplemented with 2.5 μM inhibitor (avutometinib, GDC-0623, trametiglue, trametinib, selumetinib, mirdametinib, cobimetinib, or MAP855) and 0.1% DMSO. MEK1 was prepared in the inhibitor-containing running buffer as a three-fold serial dilution series ranging from 1000 to 4.1 nM, with 0 nM included as a blank control. Samples were injected over the sensor chip at a flow rate of 30 μL min^-1^ with an association time of 120 s followed by a dissociation phase of 400 s using a multi-cycle kinetic protocol. Sensorgrams were analyzed using Biacore Evaluation Software with a 1:1 Langmuir binding model to determine the kinetic parameters.

## Data availability

All original code has been deposited on Github (https://github.com/liaulab/2026_MAPK) and is publicly available as of the date of publication. Code for BE-scan is available at https://github.com/liaulab/be-scan. The cryo-EM maps of the heterotetrameric CRAF^E393G^/MEK1 complex have been deposited in the EM Data Bank (https://www.ebi.ac.uk/emdb/) under accession code EMD-78408 and EMD-78409. The atomic model has been deposited in the Protein Data Bank (PDB) and is available at www.rcsb.org under accession code 37QE. Detailed information for the cryo-EM map and atomic model generated in this work is provided in Supplementary Table 9.

## Acknowledgements

We would like to thank members of the Liau Lab, especially J.W. Morriss and C.R. Kim for helpful discussions and comments on the manuscript. We additionally thank Dr. Andrew Aguirre and Dr. Jason Kwon for very helpful discussions and input during the conception and execution of this study and for providing a collated dataset of KRAS deep mutational scans from the literature and Jeffery Nelson for assistance with FACS. This work was supported by the National Institute of General Medical Sciences of the National Institutes of Health under award number R35GM153476. J.W. was supported by the National Institutes of Health Ruth L. Kirschstein National Research Service Award (NRSA) Individual Predoctoral Fellowship from the National Cancer Institute (F31CA294870) and previously by the Fuji Film and Harvard Therapeutics Program fellowship and the Harvard Chemical Biology Training grant (T32GM139775). H.S.K. was supported by Charles A. King Trust Postdoctoral Research Fellowship from Sara Elizabeth O’Brien Trust/Simeon J. Fortin Charitable Foundation, Bank of America Private Bank, and co-trustees. M.J.E. was supported by the National Cancer Institute of the National Institutes of Health (R35CA242461).

## Author contributions

J.W. and B.B.L. conceived the study and designed experiments; J.W performed base editor scanning; C.X.H. and J.W. conceive, designed, and performed computational analysis of base editor scanning with input from B.B.L., S.C., C.F.; J.W. and C.F. performed and analyzed validation cell-based experiments; J.W., H.S.K., and I.I. conducted preliminary base editor scanning; D.M.J performed and analyzed cryo-EM experiments, protein purification, and in vitro biochemistry experiments with input from M.J.E; J.W., C.X.H., B.B.L., D.M.J., and M.J.E. wrote and edited the paper, with inputs from all authors; B.B.L. held overall responsibility for the study.

## Competing interests

B.B.L. is a cofounder, member of the scientific advisory board and holds equity in Light Horse Therapeutics, holds equity in Aluco BioSciences, and receives financial support from Ono Pharmaceuticals Co., Ltd. The remaining authors declare no competing interests.

---

**Extended Data Fig. 1:**
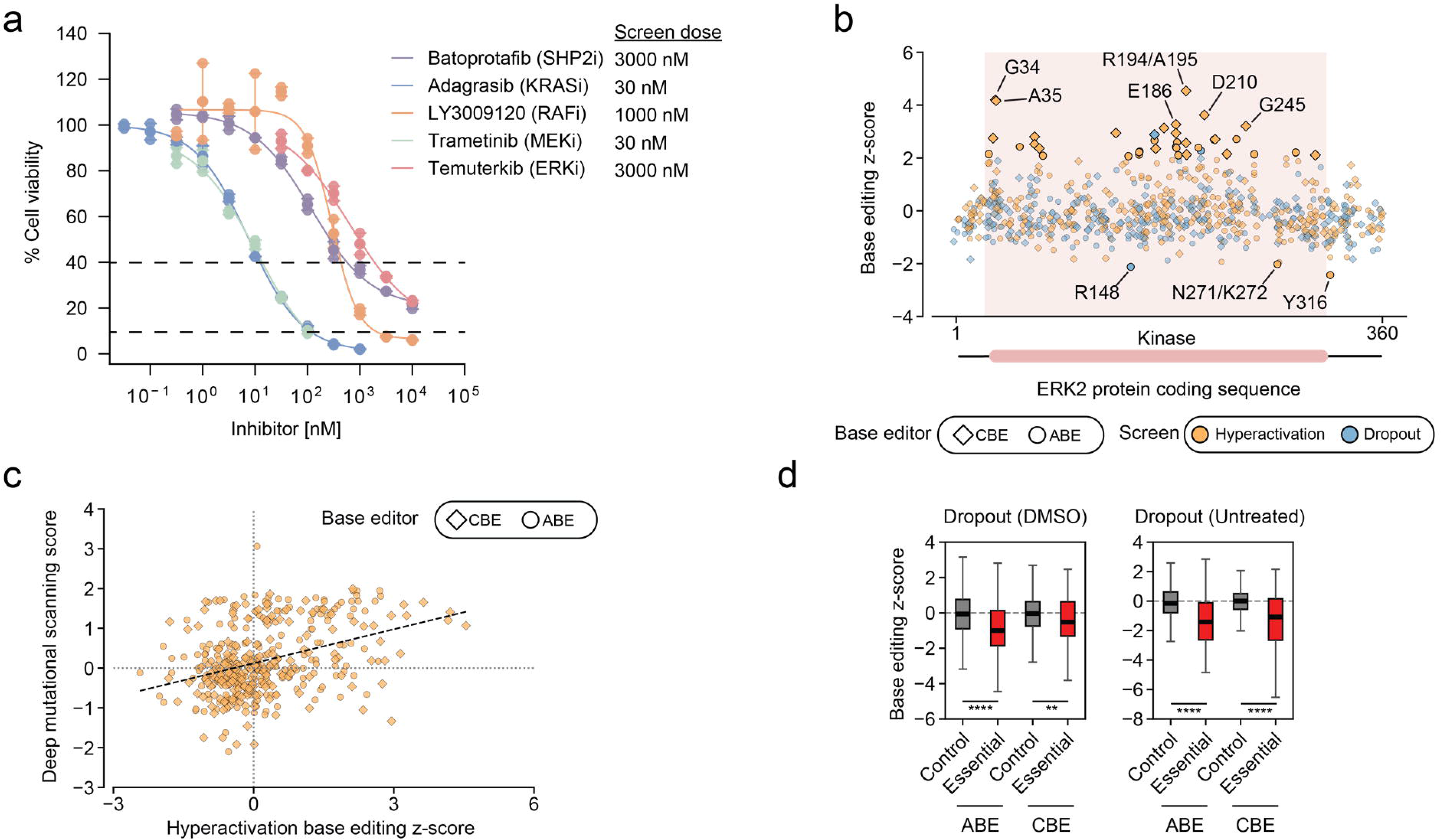
Supporting evidence for base editor scans of the MAPK pathway. **a**, Five-day growth assays (CellTiter-Glo) with five MAPK pathway inhibitors in H358 cells. Assay was performed in duplicate and normalized to DMSO control. Plots combine data collected in two separate experiments; RAFi and MEKi were assayed separate from the other inhibitors. **b**, Scatter plot showing Z-scores of sgRNAs targeting the ERK2 protein coding sequence from ABE and CBE hyperactivation positive-selection and dropout negative-selection loss-of-function data in H358-tetO-KRAS cells. Negative Z-scores indicate loss-of-function in dropout mode, while positive hits indicate loss-of-function in hyperactivation mode. Scores were calculated by comparing sgRNA abundance after 15 days between doxycycline-treated and an untreated group (hyperactivation) or the untreated group and day 0 (dropout). Solid dots indicate sgRNAs with |Z-scores| > 3. Selected sgRNAs are labeled with their predicted edited residues. **c**, Scatter plot showing correlation between ERK2 BE hyperactivation scan scores and independent ERK2 deep mutational scanning data^28^ (Pearson 0.4546, p value < 0.001). Each point in the plot is an sgRNA. A DMS score is calculated for each sgRNA by taking the average of all the DMS Z-scores for the predicted mutations of a given sgRNA. **d**, Boxplots comparing Z-scores of nontargeting sgRNAs and sgRNAs predicted to target splice site sequences of essential genes. The untreated plot shows data from the LOF screen in H358-tetO-KRAS cells, and the DMSO plot shows data from five-node inhibitor scan. P values calculated from 2-sided Student’s t-test. Significance thresholds are defined as: * p ≤ 0.05, ** p ≤ 0.01, *** p ≤ 0.001, and **** p ≤ 0.0001.

**Extended Data Fig. 2:**
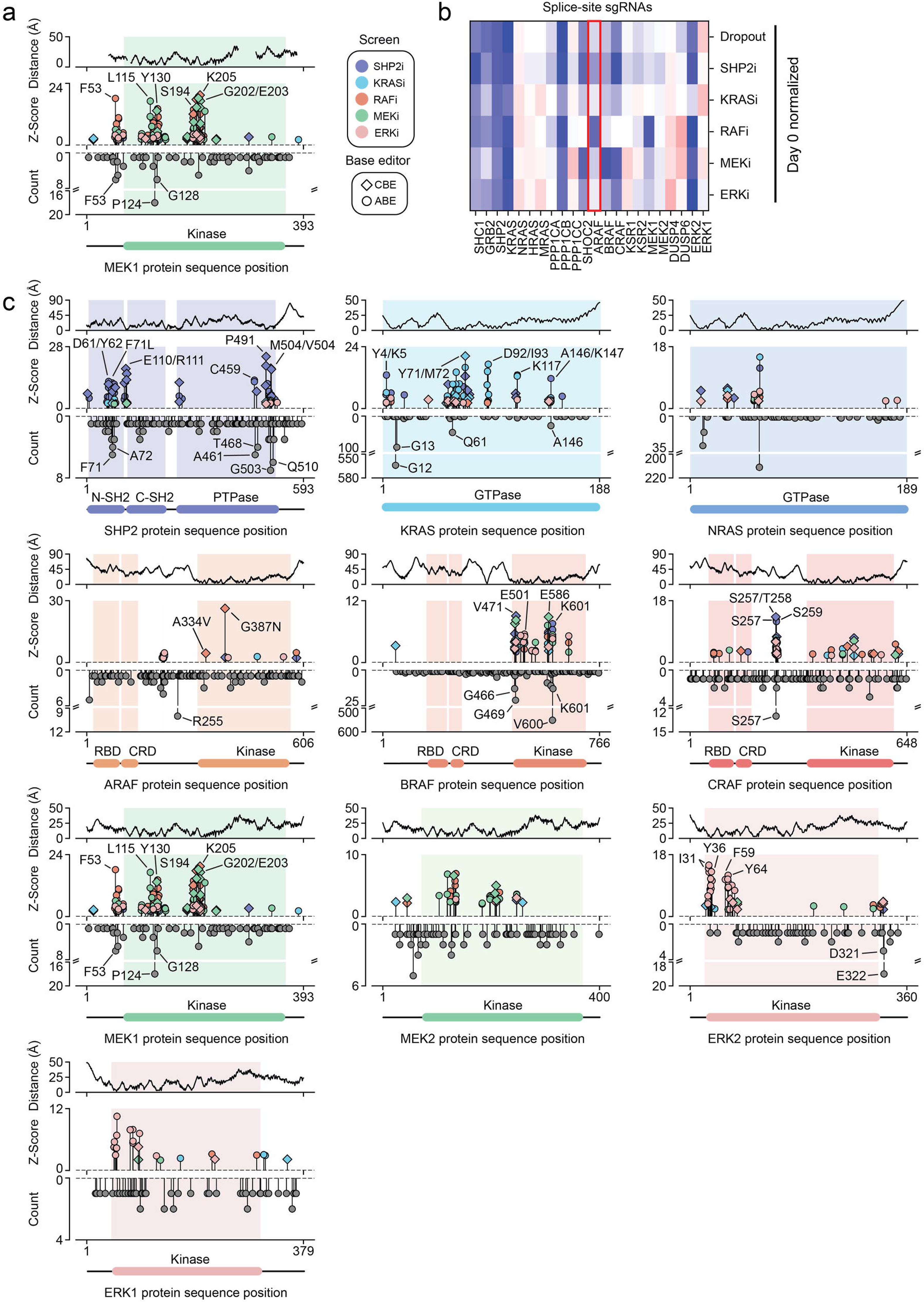
Validated inhibitor-resistant sgRNAs target clinical mutation hotspots. **a,** Plots for MEK1 showing validated resistance sgRNAs (Z-scores > 2 in validation screen) along the length of the MEK1 protein coding sequence (middle y-axis). Each axis also shows somatic mutation counts from cBioPortal (bottom y-axis) and distance of each residue from the corresponding inhibitor derived from the structure (PDB: 7JUR) (top y-axis). **b,** Splice-site and premature-stop-codon sgRNA mean Z-scores for each condition normalized to day 0 (n=5 for ABE, n=5 for CBE, all combined) as in **Fig. 2i** for all genes. **c,** Plots as in **a** but using AlphaFold structures. Inhibitors were posed by aligning known structures used in **a, Fig. 2c-e**, **Fig 3a** to AlphaFold predictions for each paralog (see Methods).

**Extended Data Fig. 3:**
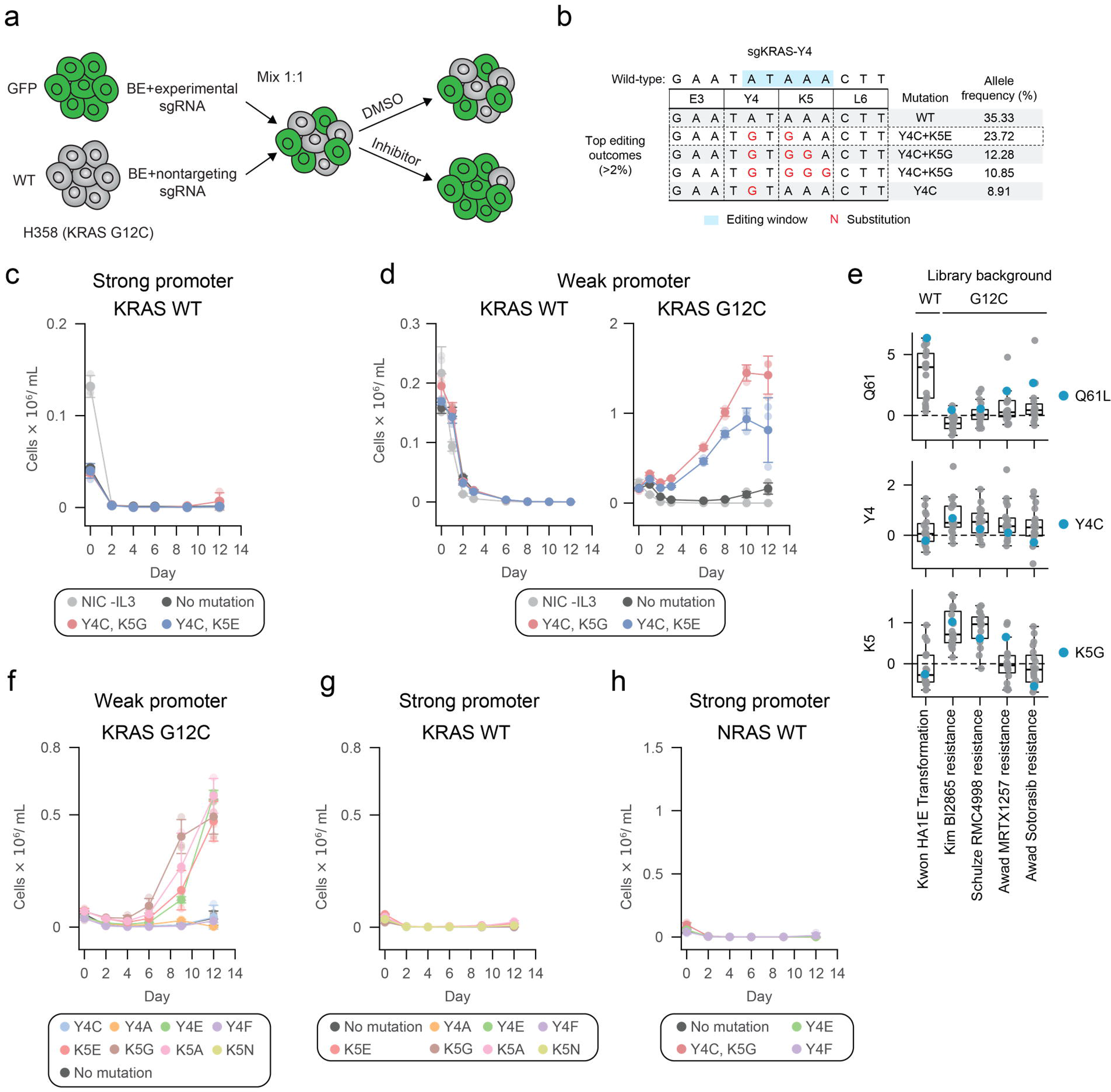
Supporting evidence for the function of KRAS residues 4 and 5. **a,** Schematic of competitive growth assay for single sgRNA validation. H358 cells (wild-type or GFP-expressing) were transduced with all-in-one base editor vectors carrying experimental or control sgRNA. After puromycin selection they are mixed 1:1, exposed to inhibitor or DMSO control, and analyzed periodically by flow cytometry. **b,** Targeted sequencing of H358 cells edited with ABE sgKRAS-Y4. **c,** Ba/F3 transformation assay measuring cell count over time with KRAS variants in wild-type background expressed from a strong promoter (EF1α). Data are presented as mean +/- standard deviation (n = 3). **d,** Ba/F3 transformation assay as in c with single mutations of KRAS but expressed from a plasmid with a weak promoter, hPGK. Left: KRAS variants in wild-type background. Right: KRAS variants in cis with G12C. **e,** Z-scores from select KRAS literature deep mutational scans.^15,22,39,40^ Mutations at Y4 and K5 are associated with activation or drug resistance but only in combination with G12C. **f,** Ba/F3 transformation assay as in d with single mutations in KRAS in cis with G12C, expressed from a weak promoter, hPGK. **g,** Ba/F3 transformation assay as in c with single mutations in KRAS*^WT^* background. **h,** Ba/F3 transformation assay as in c with single mutations in NRAS wild-type background.

**Extended Data Fig. 4:**
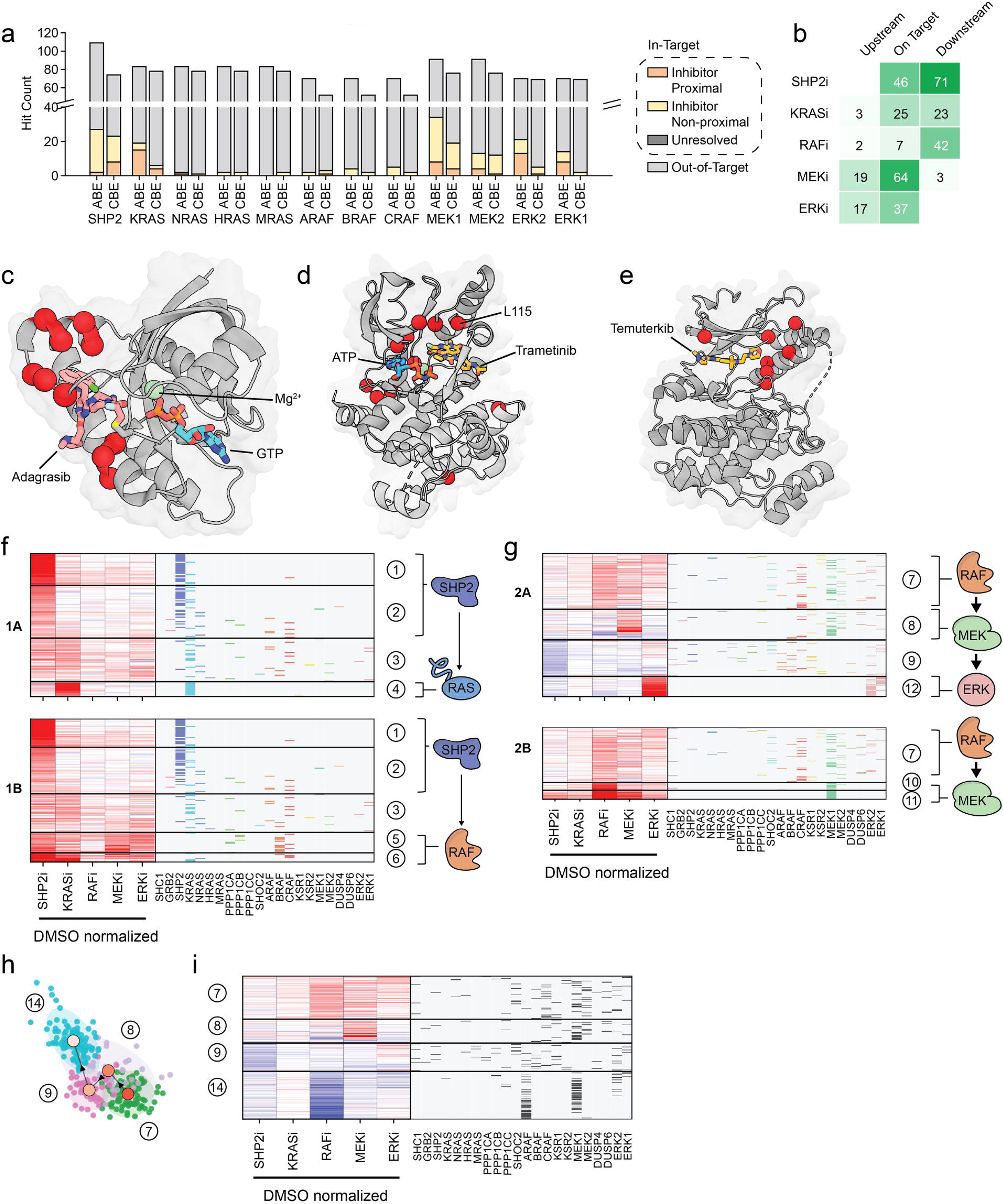
Supporting evidence for clustering and trajectory analyses. **a,** Bar plot summarizing locations of validated resistance sgRNAs using AlphaFold structures, similar to **Fig. 4a**. PDB structures used in **Fig. 4a** were used to align inhibitors and AlphaFold models. **b,** Heatmap of validated hit sgRNA counts as in **Fig. 4b** categorized by whether an sgRNA is upstream, on target, or downstream of the relevant node for each inhibitor. **c,** Predicted residues edited by KRASi-specific cluster (cluster 4) indicated on KRAS-adagrasib structure (PDB: 9O0R). **d,** MEKi-specific cluster (cluster 8) indicated on MEK1-trametinib-ATP structure (PDB: 7JUR). **e,** ERKi-specific cluster (cluster 12) indicated on ERK2-Temuterkib structure (PDB: 6RQ4). **f-g,** Heatmap excerpts as in **Fig. 4c** for each sub-trajectory of the MST analysis for trajectories 1A and 1B **f** and 2A and 2B **g** each with a cartoon indicating their dominant genes. **h-i,** Specific sub-trajectory 2C summarized in PCA space **h** and as a heatmap **i** as in **Fig. 4d and 4f-g** respectively.

**Extended Data Fig. 5:**
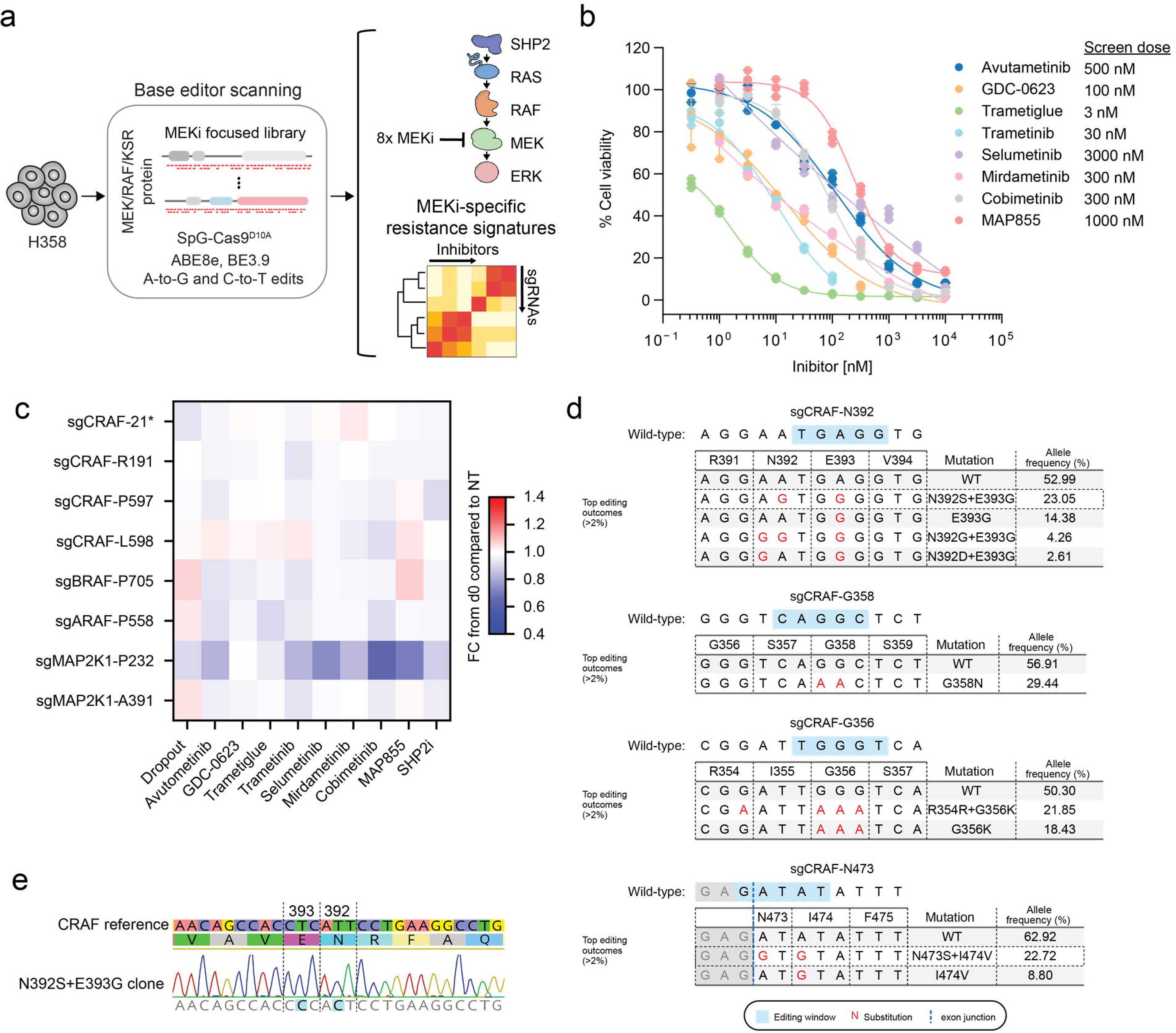
Supporting evidence for MEK inhibitor-focused BE scanning. **a,** Schematic of MEK inhibitor-focused BE scanning. **b,** Five-day growth assays (CellTiter-Glo) with 8 MEK inhibitors. Assay was performed with duplicate or triplicate wells and normalized to DMSO control. Plots combine data collected from four separate experiments, as inhibitors were not all assessed on the same day. **c,** Single sgRNA validation (competitive growth assay) for select sgRNAs run with a single endpoint at 12 days. Heatmap shows mean of 3 replicates normalized to day 0 and then normalized to a nontargeting control. See methods for inhibitor concentrations. Color indicates fold change of GFP from day 0 compared to NT. **d,** Targeted sequencing of base edited sites from single sgRNAs in **Fig. 5d**. Genotyping was performed on cell mixtures from DMSO-treated cells after twelve days of competitive growth. **e,** Targeted Sanger sequencing of clonal base edited H358 cells carrying E393G+N392S.

**Extended Data Fig. 6:**
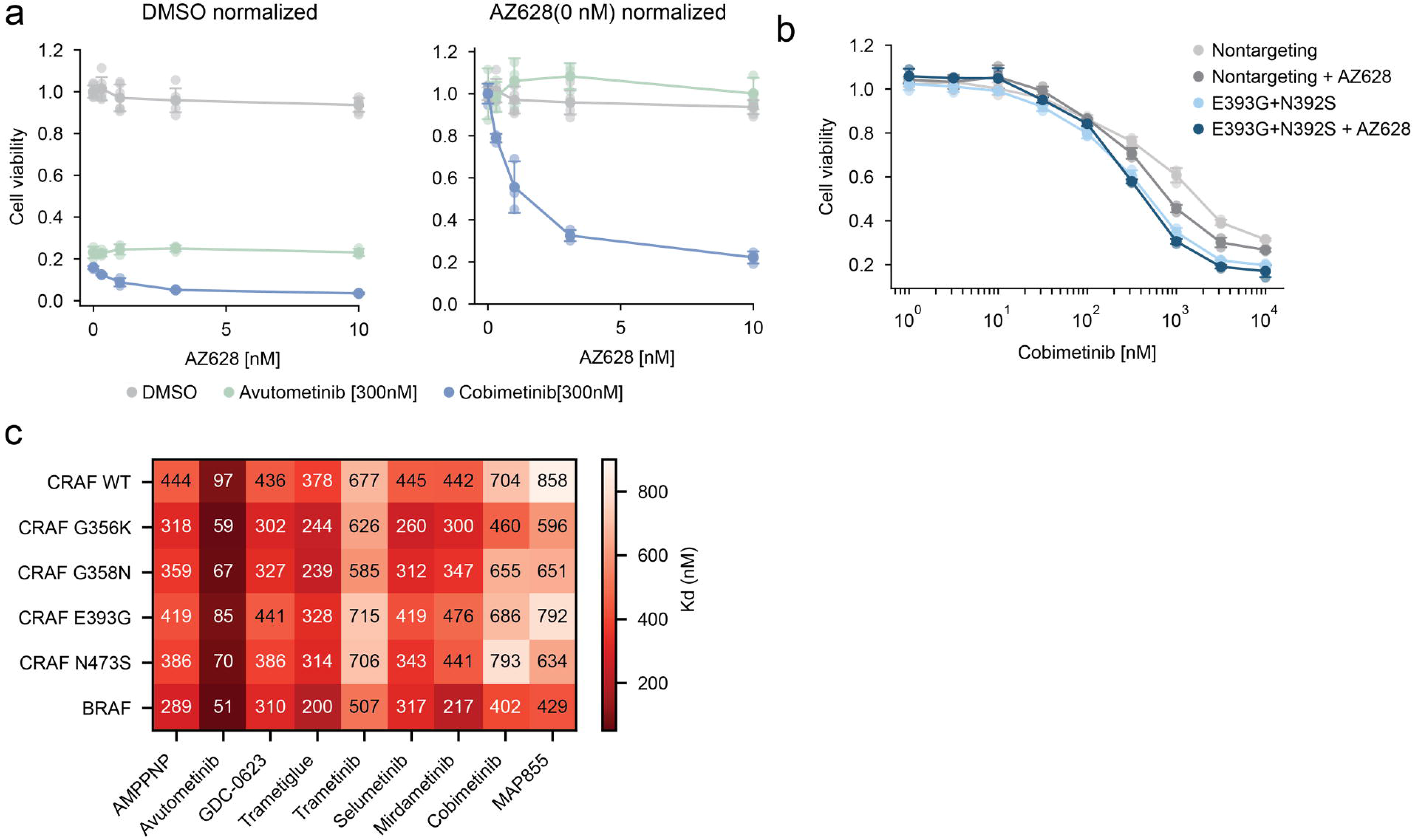
Cellular and biochemical characterization of bidirectional CRAF mutations. **a,** Five-day growth assay (CellTiter-Glo) showing dose-response of CRAF-biased inhibitor AZ628 in combination with DMSO, avutometinib (300nM), or cobimetinib (300nM). The left plot shows viability normalized to DMSO-treated cells; the right plot shows the same data with each background treatment (DMSO, avutometinib, or cobimetinib) independently normalized to its own 0 nM AZ628 value. Performed in technical triplicate. **b,** Cobimetinib dose response with homozygous CRAF E393G+N392S edited clone or clonal cell line carrying a non-targeting sgRNA with and without AZ628 (3nM). **c,** SPR using immobilized CRAF kinase domain with indicated mutations or BRAF kinase domain. Affinity for MEK1 (S218A/S222A mutant) measured in the presence of various MEK inhibitors (2.5μM). AMPPNP was included in all conditions (5μM).

**Extended Data Fig. 7:**
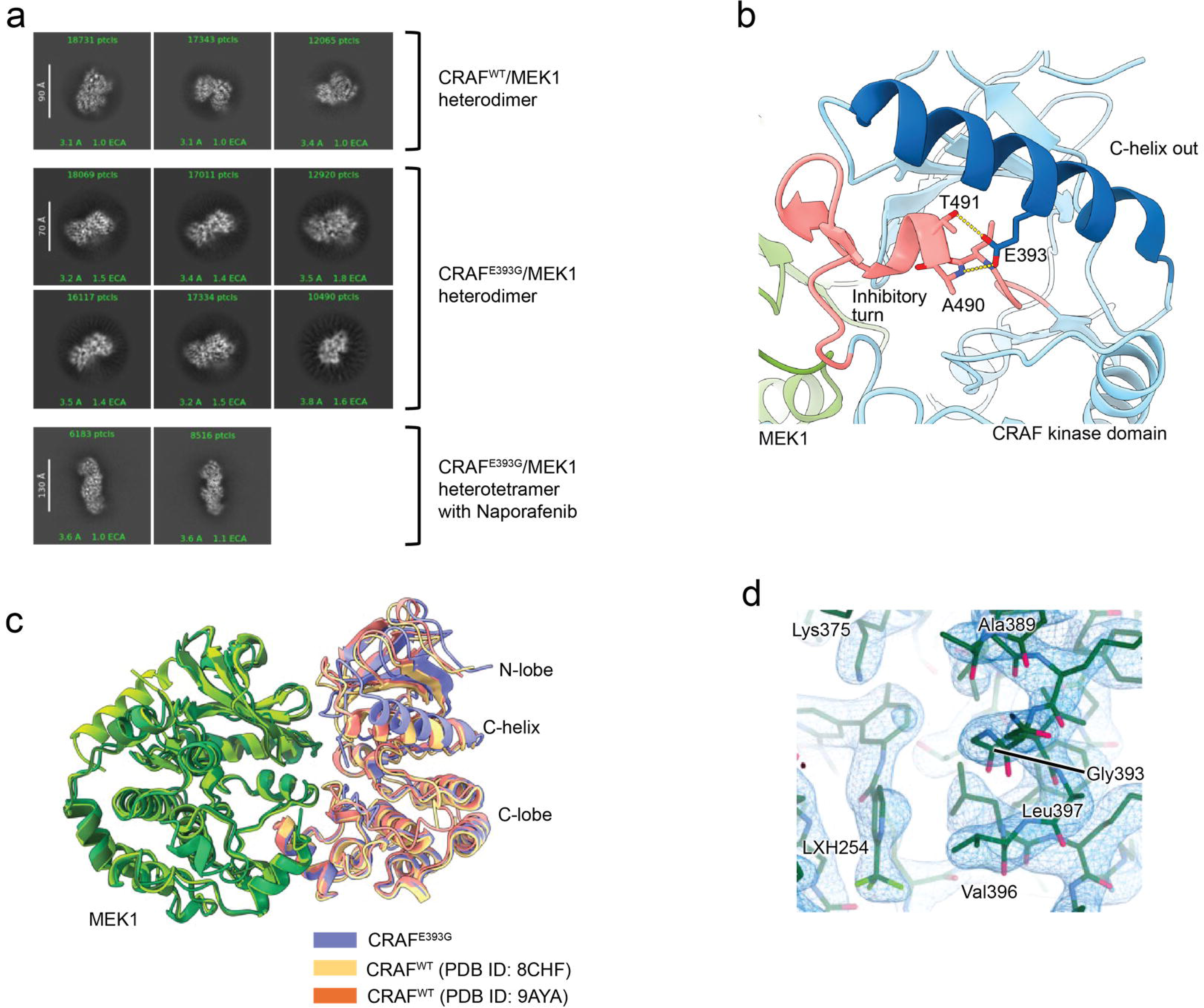
Cryo-EM structure analysis of the CRAF^E393G^/MEK1 complex. **a**, Representative 2D class images of the heterodimeric CRAF^WT^/MEK1 with an ATP analog (from the dataset used for PDB ID: 9MMQ), the heterodimeric CRAF^E393G^/MEK1 with an ATP analog (dataset not deposited), and the heterotetrameric CRAF^E393G^/MEK1 with naporafenib. The scale bar (Å), particle number, and estimated resolution are indicated on the left, top, and bottom, respectively. **b**, Structural analysis of CRAF^WT^/MEK1 complex. The CRAF^WT^/MEK1 complex in the inactive conformation (PDB ID: 9MMQ) is shown in ribbon representation. CRAF^WT^ and MEK1 are colored in light blue and green, respectively. The C-helix and inhibitory turn in CRAF^WT^ are highlighted in dark blue and salmon, respectively. **c**, Structural comparison of the CRAF^E393G^/MEK1, CRAF^WT^/MEK1/14-3-3 (C-helix in), and CRAF^WT^/MEK1 (C-helix in) complexes. The CRAF/MEK1 dimer from the heterotetrameric CRAF^E393G^/MEK1 complex was superposed with that from the heterotetrameric CRAF^WT^/MEK1/14-3-3 (PDB ID: 8CHF) and from the heterotetrameric CRAF^WT^/MEK1 (PDB ID: 9AYA). MEK1 is colored green, and CRAF is colored as indicated in the color key below. Note that the C-helix of CRAF^E393G^ is shifted outward related to CRAF^WT^. **d**, Cryo-EM density surrounding the CRAF^E393G^ mutation site. The map is shown as a blue mesh.

**Extended Data Fig. 8:**
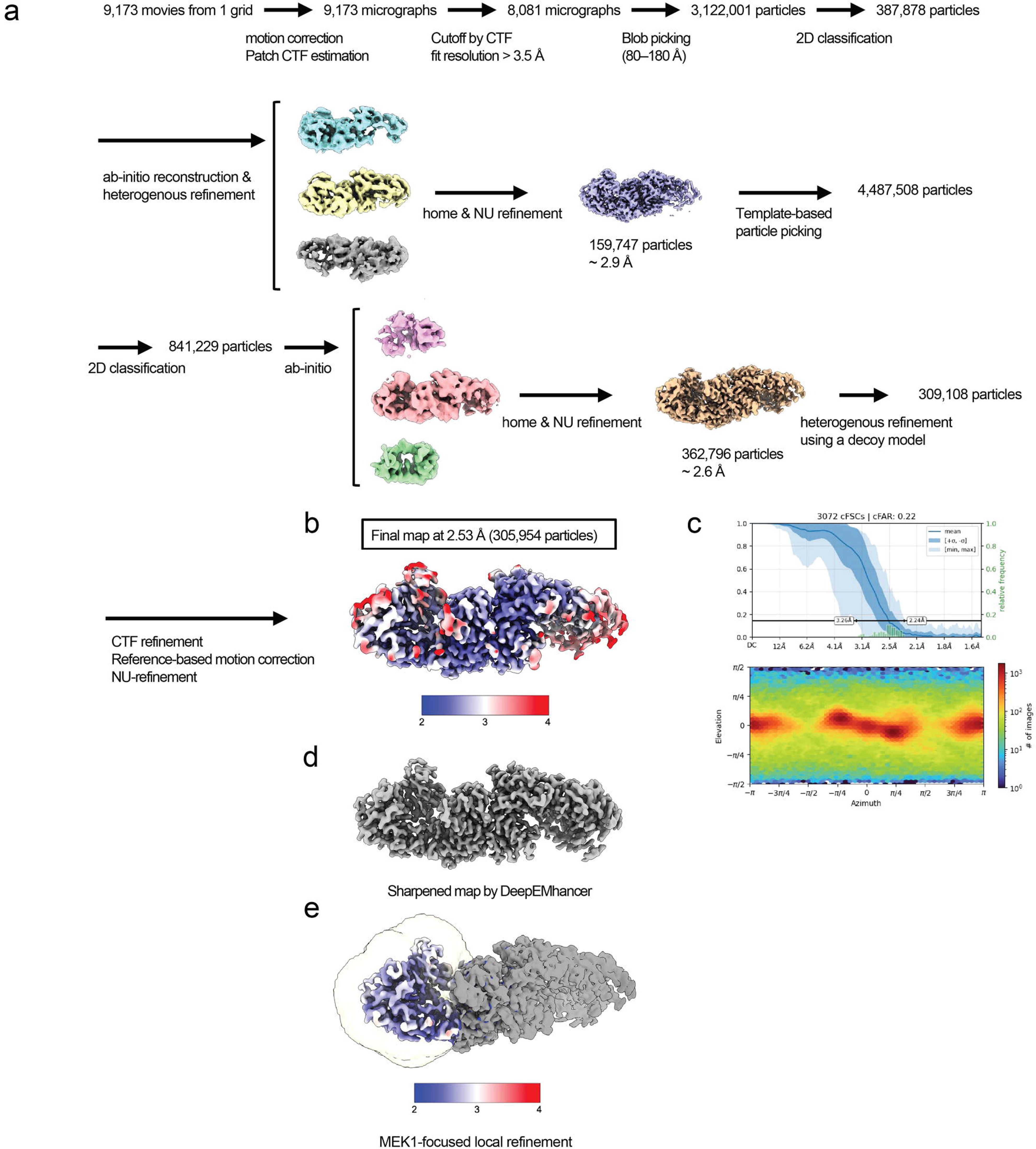
Overall cryo-EM data processing for the heterotetrameric CRAF^E393G^/MEK1 complex. **a**, Cryo-EM data processing workflow for the heterotetrameric CRAF^E393G^/MEK1 complex. **b**, Final cryo-EM density map colored by local resolution. The overall resolution was estimated from gold-standard Fourier shell correlation (GSFSC) curve calculated in CryoSPARC. **c**, Corresponding conical Fourier shell correlation (cFSC) plot and direction distribution plot calculated in CryoSPARC. **d**, Cryo-EM density map sharpened by DeepEMhancer. This sharpened map was used for structural analysis shown in Fig. 6d. **e**, MEK1-focused local refinement. A yellow mask was applied during local refinement, resulting in improved map quality and local resolution around MEK1.

## Notes

https://liaulab.github.io/2026_MAPK/

