## Supplementary Table 9 for "Pathway-wide base editing charts chemical-genetic interactions in MAPK signaling"

**Supplementary Table 9 | Cryo-EM data collection, refinement and validation statistics for the heterotetrameric CRAF^E393G^/MEK1 complex.**

| **Data collection and processing** |  |
| --- | --- |
| Magnification | 165,000 x |
| Voltage (kV) | 300 |
| Electron exposure (e/Å^2^) | 49.61 |
| Defocus range (μm) | − 0.8 – −1.8 |
| Pixel size | 0.74 |
| Symmetry imposed | C1 |
| Number of micrographs | 9,185 |
| **Refinement** | (PDB xxxx)  (EMD-xxxxx) |
| Final particle images (no.) | 305,954 |
| Map resolution (Å) |  |
| 0.143 FSC threshold | 2.5 (masked), 3.1 (no mask) |
| Initial model used (PDB code) | 9MMQ |
| Model composition |  |
| Chains | 4 |
| Non-hydrogen atoms | 8811 |
| Protein residues | 1088 |
| Ligands | 6 |
| Metals | 1 |
| *B* factors (Å^2^) |  |
| Proteins (min/max/average) | 65.36/267.25/133.19 |
| Ligands (min/max/average) | 72.20/239.54/136.64 |
| R.m.s. deviations |  |
| Bond length (Å) | 0.005 |
| Bond angles (°) | 0.692 |
| Validation |  |
| MolProbity score | 1.65 |
| Clash score | 7.44 |
| Poor rotamers (%) | 0.53 |
| Ramachandran plot |  |
| Favored (%) | 96.36 |
| Allowed (%) | 3.64 |
| Outliers (%) | 0.0 |
| Model vs. Data |  |
| CC (mask) | 0.83 |
| CC (box) | 0.78 |
| CC (peaks) | 0.70 |
| CC (volume) | 0.83 |
| Mean CC for ligands | 0.72 |
